# Snip1 and PRC2 coordinate intrinsic apoptosis, cell division, and neurogenesis in the developing brain

**DOI:** 10.1101/2022.04.27.489801

**Authors:** Yurika Matsui, Mohamed Nadhir Djekidel, Katherine Lindsay, Parimal Samir, Nina Connolly, Hongfeng Chen, Yiping Fan, Beisi Xu, Jamy C. Peng

## Abstract

Brain development requires the intricate balance between division, death, and differentiation of neural progenitor cells (NPCs). Here, we report the discovery of Snip1 as a key regulator of these NPC phases. The conditional deletion of *Snip1* in the mouse embryonic brain causes dysplasia with robust induction of caspase 9-dependent apoptosis. In NPCs, Snip1 suppresses the genetic programs of apoptosis and developmental signaling pathways and promotes the genetic programs of cell cycle, neurogenesis, and cortical development. Mechanistically, Snip1 binds to the Polycomb complex PRC2, co-occupies gene targets with PRC2, and regulates H3K27 marks. Deletion of PRC2 is sufficient to reduce apoptosis and brain dysplasia and partially restore genetic programs and tissue development in the Snip1-depleted brain. Our findings suggest that Snip1 exerts loci-dependent regulation of PRC2 and H3K27me3 to toggle between cell fates in the developing brain.

## INTRODUCTION

Neural progenitor cells (NPCs) are self-renewing and multipotent cells that give rise to neurons, oligodendrocytes, and astrocytes in the central nervous system (CNS). The precise size and structural organization of the brain is achieved by the exquisite balance between division, death, and differentiation of NPCs. Cell death is prominent in the normal developing brain, especially in the proliferative zones of the cortex where NPCs reside ^1-4^. However, how NPCs are developmentally programmed to toggle between survival and death is poorly understood.

The cell-intrinsic control of NPCs is mediated through epigenetic mechanisms, and a key epigenetic modifier is Polycomb repressive complex 2 (PRC2). PRC2 deposits H3K27 trimethylation (H3K27me3), mediates chromatin compaction, and suppresses RNA polymerase II–dependent transcription ^5-8^. Genetic deletion of PRC2 subunits causes embryonic lethality with gastrulation failure ^8-11^. PRC2 plays essential roles throughout CNS development, such as neural tube closure, expansion of NPCs, and cell fate specification ^12-14^. Additionally, PRC2-mediated H3K27me3 is correlated with neuroprotection ^5,15,16^. Aberrant increases of H3K27me3 and higher stability of the PRC2 enzymatic subunit Ezh2 are observed in ataxia-telangiectasia ^15^, whereas H3K27me3 reduction is linked to Parkinson’s disease and Huntington’s disease ^5,16^. PRC2 and H3K27me3 suppress gene targets involved in neurodegeneration in the adult brain ^5,15^. In these disease models, the dysregulation of PRC2 and H3K27me3 leads to neural loss.

Snip1 was originally discovered by a yeast two-hybrid screen for interactors of SMAD proteins ^17^. The N-terminus of SNIP1 binds to histone acetyltransferases p300/CBP at their CH1 domain to disrupt the SMAD4–p300/CBP complex formation and dampen BMP/TGFβ signaling ^18-21^. SNIP1 can also dampen NFκB signaling by disrupting RELA/p65-p300/CBP complex formation ^22-26^. Besides its role in TGFβ or NFκB signaling, SNIP1 acts on oncogenes or tumor suppressors to promote cell transformation and cell cycle. In many cancer cells, SNIP1 can increase the stability of c-MYC and its complex with to p300/CBP ^27^, disrupt the RB-HDAC1 complex ^28^, or promote cyclin D1 expression ^29,30^. Ectopic expression of Snip1 induces defective patterning in the *Xenopus* embryos ^17^. Global knockout (KO) of Snip1 in zebrafish embryos causes reduction in GABAergic and glutamatergic neurons ^31^.

Using Snip1 conditional KO in NPCs, we revealed that Snip1 suppresses intrinsic apoptosis and promotes cell cycle progression and cortical development. Mechanistically, Snip1 binds to PRC2 and co-occupies their gene targets to regulate H3K27 marks and gene expression. The depletion of a PRC2 subunit Eed partially restores gene expression, NPC function, and development in the Snip1-KO brain. Our study reveals a novel epigenetic pathway that is crucial for regulating the properties of NPCs in the developing brain.

## RESULTS

### Snip1 is required for the survival of NPCs in the murine embryonic brain

We examined the expression of Snip1 in the murine embryonic brain by RNAscope ^32^. At embryonic day E11.5 and E13.5, *Snip1* transcripts were expressed in nearly all cells and robustly expressed in the neuroepithelia lining the ventricles, where NPCs reside (Supp Fig 1a-b). To study Snip1 in the murine embryonic brain, we used *Nestin* (*Nes*)::*Cre* to conditionally deplete Snip1 in NPCs, hereafter referred to as *Snip1*^*Nes*^*-*KO. *Nes*::Cre is expressed in NPCs to recombine the flox sites and excise exon 2 of *Snip1* (Fig 1a, Supp Fig 1c-d, Supp Fig 2a-d). By E15, *Snip1*^*Nes*^*-KO* embryos displayed severe thinning of brain tissues and dysplasia with 100% penetrance (Fig 1b-c, p <0.0001 by Fisher’s exact test). To understand the cellular underpinnings of the brain dysplasia in *Snip1*^*Nes*^*-*KO, we examined cell proliferation and apoptosis of NPCs. NPCs were identified by their expression of the neural stem cell marker Sox2. To identify proliferating cells *in vivo*, we injected BrdU into pregnant dams and/or detected the proliferative marker Ki67. Quantification of these markers in neuroepithelia did not reveal significant difference in proliferative NPCs in sibling control and *Snip1*^*Nes*^*-*KO embryos (Supp Fig 2e-h).

**Fig. 1.**
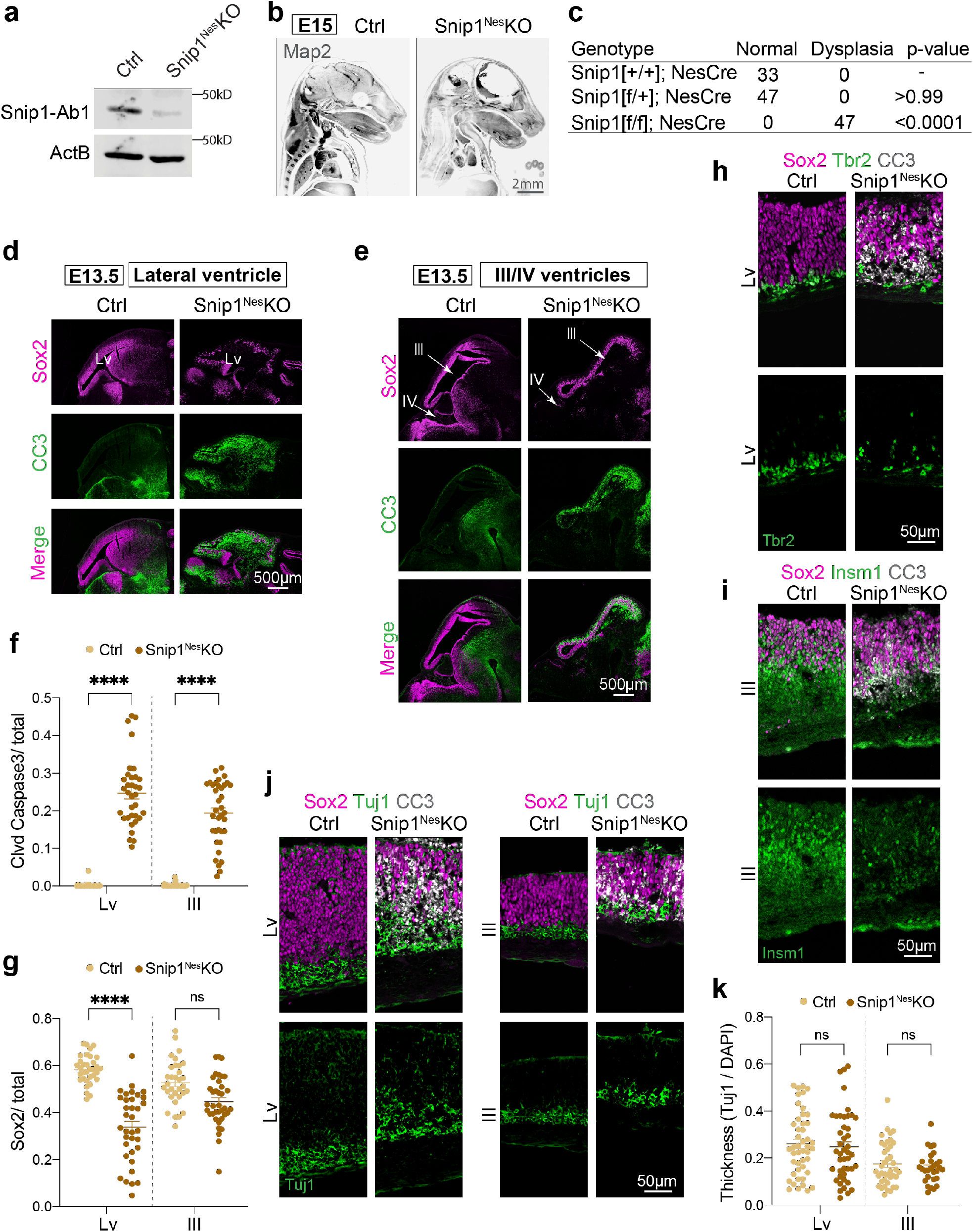
Depletion of *Snip1* in NPCs causes brain dysplasia in mouse embryos. **a** WB of control and *Snip1*^*Nes*^-KO NPCs at E13.5. Snip1-Ab1; anti-Snip1 antibody from ProteinTech. **b** IF of Map2 in control and *Snip1*^*Nes*^-KO embryos at E15. Embryos were cleared by the iDISCO method and imaged with light sheet microscopy. Bar, 2 mm. **c** Penetrance of brain dysplasia in E13.5 embryos. Brain dysplasia was determined by the thinning of the brain tissue. Statistical significance was calculated by Fisher’s exact test. **d-e** IF of Sox2 and cleaved caspase 3 (CC3) in sagittal cryosections of the E13.5 brain. Germinal zones around (**d**) lateral ventricle (Lv, forebrain) and (**e**) 3^rd^ (midbrain) /4^th^ (hindbrain) ventricles were examined. Bar, 500 μm. **f-g** Quantification of CC3-positive and Sox2-positive cells in the neuroepithelial lining of the ventricles of control and *Snip1*^*Nes*^-KO embryos at E13.5. DAPI staining was used to count the total number of cells. Each data point represents one image. N=8 embryos for control and n=7 for *Snip1*^*Nes*^-KO. Data are presented as mean ± SEM, and two-way ANOVA was used for statistical analysis. ns = not statistically significant; ****p <0.0001. **h-j** IF of Sox2 and CC3 overlayed with neural lineage markers, Tbr2, Insm1, or Tuj1 of the E13.5 brain. Bar, 50 μm. **k** Thickness of the Tuj1-positive region relative to the entire cortical thickness. Each data point represents one image. N=11 embryos for control and n=7-11 for *Snip1*^*Nes*^-KO. Data are presented as mean ± SEM, and two-way ANOVA was used for statistical analysis. ns = not statistically significant.

We next probed apoptosis of NPCs by IF of cleaved (cl)-caspase 3. At E13.5, all ventricles of *Snip1*^*Nes*^*-*KO displayed strong induction of cl-caspase 3, accompanied by loss of NPCs (Fig 1d-g). Given that cl-caspase 3 signals were enriched in the subventricular zones (SVZs) of neuroepithelia, where most intermediate progenitors reside, we examined the markers of intermediate progenitors Tbr2 and Insm1. Compared with the sibling control, Tbr2 and Insm1 were markedly reduced in *Snip1*^*Nes*^*-*KO neuroepithelia (Fig 1h-i). By E13.5, the embryonic brain undergoes neurogenesis. Therefore, we examined the immature neuron marker Tuj1. The relative thickness of the Tuj1-positive region did not significantly differ between the *Snip1*^*Nes*^*-*KO and control at E13.5 (Fig 1j-k). These data suggest that Snip1 suppresses apoptosis in NPCs and intermediate progenitors in the developing brain.

To eliminate the possibility that the observed apoptosis phenotype is not due to *Nes*::Cre, we used *Emx1*::Cre to conditionally deplete Snip1 in NPCs of the dorsal telencephalon ^33^. The *Snip1*^*Emx1*^*-* KO embryos showed strong induction of apoptosis, loss of Tbr2-positive intermediate progenitors, and dysplasia of the forebrain (Supp Fig 2i-k). These findings support that in the developing murine brain, apoptosis induction and dysplasia are independent of *Nes*::Cre and specific to Snip1 depletion. These data further support our conclusion that Snip1 is required for an anti-apoptotic and pro-survival mechanism in NPCs and intermediate progenitors.

### Snip1 promotes cortical development and suppresses the intrinsic apoptosis program

To study the temporal dynamics of apoptosis in the *Snip1*^*Nes*^*-*KO brain, we examined cl-caspase 3 in the E11.5 embryos. By E11.5, cl-caspase 3 signals were detected throughout the *Snip1*^*Nes*^*-*KO brain (Supp Fig 3a-d). As *Nes*::Cre is turned on by E10.5 ^34^, Snip1 depletion likely induces apoptosis within 24 hours. The quantification of Sox2-positive cells in lateral, third, and fourth ventricles revealed that only NPCs in the fourth ventricle begin to be depleted by E11.5 (Supp Fig 3e). Hereafter, we focused our analyses of lateral and third ventricles to study apoptosis control by Snip1.

To shed light on the molecular underpinnings of the defective *Snip1*^*Nes*^*-*KO NPCs, we performed RNA-sequencing (RNA-seq) of Sox2-positive NPCs sorted from E13.5 *Snip1*^*Nes*^*-*KO and sibling controls (Fig 2a, Supp Fig 4a). We analyzed genes with count per million (CPM) values >1 in either control or *Snip1*^*Nes*^*-*KO NPCs. Using the criteria of false discovery rate (FDR) <0.05 to compare 4-replicate datasets each from control and *Snip1*^*Nes*^*-KO* NPCs, we identified 1,210 upregulated genes and 1,621 downregulated genes in *Snip1*^*Nes*^-KO (Fig 2b, Supp Fig 4b). Gene set enrichment analysis (GSEA) revealed that upregulated genes in *Snip1*^*Nes*^*-*KO NPCs were enriched in functions related to apoptosis, H3K27me3 or bivalent promoters in NPCs and the brain, midbrain markers, spliceosomal small nuclear ribonucleoprotein particles, and signaling pathways involving TNF, IGF, TGFβ, and Hedgehog (Fig 2c). Downregulated genes in *Snip1*^*Nes*^*-*KO NPCs were enriched in functions related to forebrain and cortex development, CNS neuron differentiation, chromosome segregation, NPC proliferation, axonogenesis, replication fork, and signaling pathways involving TLR and Rho (Fig 2d).

**Fig. 2.**
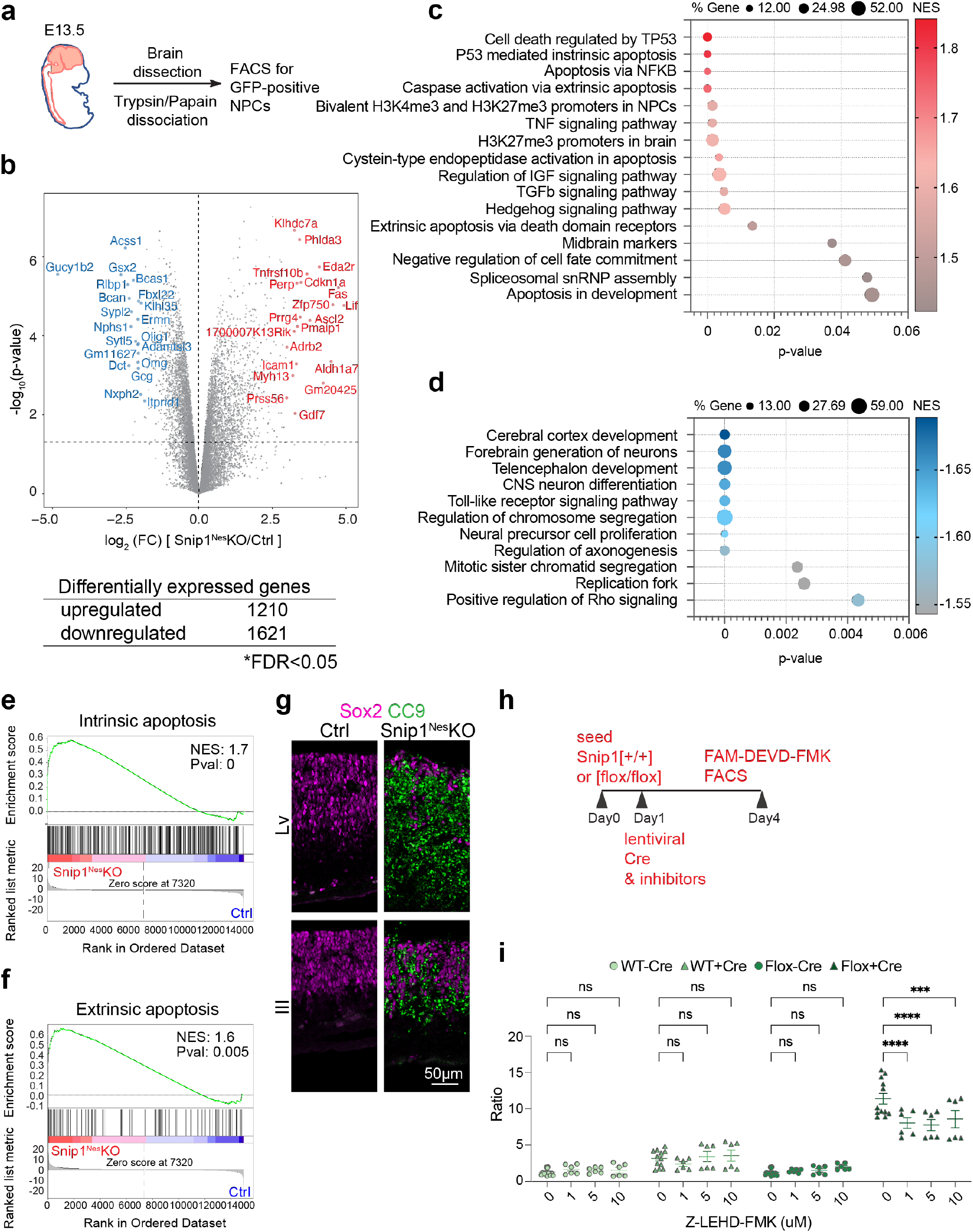
Snip1 suppresses genes involved in apoptosis and signal transduction and promotes genes for brain development. **a** Schematic of the brain NPC collection. **b** Volcano plot and the number of differentially expressed genes between the control and *Snip1*^*Nes*^-KO NPCs. The table shows the number of genes that passed the cutoff of FDR <0.05. FDR was calculated by the Benjamini & Hochberg method. **c-d** Bubble plots of the enriched gene sets in (**c**) upregulated genes and (**d**) downregulated genes in *Snip1*^*Nes*^-KO vs. control NPCs. Differentially expressed genes were first ranked by their fold-change, p-value, and expression level before Gene Set Enrichment Analysis (GSEA) was performed. P-values were calculated by one-tailed Kolmogorov–Smirnov test. **e-f** Representative GSEA of upregulated genes in *Snip1*^*Nes*^-KO vs. control NPCs. Upregulated genes were enriched in gene sets related to both (**e**) intrinsic and (**f**) extrinsic apoptosis. Differentially expressed genes were first ranked by their fold-change and p-value before GSEA was performed. **g** IF of cleaved caspase 9 (CC9) overlayed with Sox2 in sagittal cryosections of the E13.5 brain. Bar, 50 μm. **h** Schematic of transduction with mCherry-Cre lentivirus and treatment with inhibitors in Snip1[+/+] and Snip1[flox/flox] NPCs. **i** The percentage of cells with active caspase 3 quantified by FACS. Caspase 9 inhibitor (Z-LEHD-FMK) was added at different concentrations along with mCherry-Cre lentivirus. Data are presented as mean ± SEM, and two-way ANOVA was used for statistical analysis. ns = not statistically significant; ***p <0.001; ****p <0.0001.

As intrinsic and extrinsic apoptotic signatures were enriched in the upregulated genes in *Snip1*^*Nes*^*-*KO NPCs (Fig 2c, e-f), we examined the activation of these two pathways. IF showed little signal of cl-caspase 8, an effector of the extrinsic apoptosis pathway ^35,36^; Supp Fig 4c) but strong signals of cl-caspase 9, an effector of the intrinsic apoptosis pathway ^37-39^, throughout the neuroepithelia of ventricles (Fig 2g). Additionally, quantification of cl-caspase 3 by FACS of FAM-DEVD-FMK staining ^40,41^ show that whereas inhibition of caspase 8 modestly altered apoptosis, inhibition of caspase 9 robustly reduced apoptosis in the *Snip1*^*Nes*^*-*KO NPCs (Fig 2h-i, Supp Fig 4d). For identifying a cause of increased apoptosis, we did not detect an increase in DNA damage (Supp Fig 4e-f) but detected strong increases of p53 signals in Sox2-positive NPCs (Supp Fig 4g). These data suggest that the *Snip1*^*Nes*^*-*KO embryo displayed dysregulated control of p53-mediated intrinsic apoptosis in NPCs. We propose that Snip1 primarily suppresses the intrinsic apoptosis as part of a neurodevelopmental program.

GSEA results showed that downregulated genes in the *Snip1*^*Nes*^*-*KO NPCs were enriched in forebrain developmental programs. Although the *Snip1*^*Nes*^*-*KO forebrain tissues displayed severe thinning as a consequence of apoptosis, the forebrain marker Foxg1 and the mid/hindbrain marker Otx2 were similarly detected between the control and *Snip1*^*Nes*^*-*KO brains (Supp Fig 4h). These data suggest that Snip1 depletion did not alter forebrain specification.

Other downregulated genes in the *Snip1*^*Nes*^*-*KO NPCs were involved in the control of self-renewal (Fig 2d). Characterization of the *Snip1*^*Nes*^*-*KO NPCs *in vitro* showed that, compared with control NPCs, cultured *Snip1*^*Nes*^*-*KO NPCs had reduced Sox2 expression (Supp Fig 5a). By allowing NPCs to form neurospheres in suspension and through serial passages, we observed that neurosphere number and cross-section area were significantly lower in *Snip1*^*Nes*^*-*KO compared with control (Supp Fig 5b-d). Overexpressing human SNIP1 (85% identity) was sufficient to rescue self-renewal in cultured *Snip1*^*Nes*^*-* KO NPCs (Supp Fig 5e-f), suggesting functional conservation of Snip1. These findings suggest that Snip1 is required for the self-renewal in NPCs.

We performed RNA-seq of Sox2-negative differentiated cells sorted from E13.5 *Snip1*^*Nes*^*-*KO and sibling control brains. Using the criteria of fold-change >2 and p <0.05 to compare 2-replicate datasets each from control and *Snip1*^*Nes*^*-KO*, we identified 658 upregulated genes and 150 downregulated genes in *Snip1*^*Nes*^-KO (Supp Fig 5g). GSEA revealed that upregulated genes in *Snip1*^*Nes*^*-*KO cells were enriched in functions related to apoptotic clearance, neuron specification and differentiation, midbrain markers, and known high-CpG-density promoters occupied by bivalent marks(H3K27me3 and H3K4me3) in NPCs ^42^ (Supp Fig 5h). Downregulated genes in *Snip1*^*Nes*^*-*KO cells were enriched in functions related to spliceosome, translation and ribosome, nucleosome organization, and apoptosis via p21 but not p53 (Supp Fig 5i). These results implicate that Snip1 regulates different functional gene sets in NPCs compared to differentiated cells in the developing brain.

### Snip1 directly regulates genetic programs including intrinsic apoptosis, cell cycle, and cortical development

To determine whether Snip1 proteins directly bind gene loci to regulate their expression, we first profiled the genome-wide distribution of Snip1 by CUT&RUN ^43^ in *Snip1*^*Nes*^-KO and control NPCs. Using SICER ^44^ and MACS2 ^45^ with FDR <0.05 to compare 2 datasets each from control and *Snip1*^*Nes*^*-*KO, we identified 23,188 Snip1-bound regions in control NPCs and only 4,187 regions in *Snip1*^*Nes*^*-*KO NPCs (Supp Fig 6a-c). Heatmaps of the 23,188 sites showed drastic reduction of Snip1 CUT&RUN signals in *Snip1*^*Ne*^-KO (Fig 3a). These differences suggest the high specificity of Snip1 CUT&RUN in NPCs. Approximately 50% of Snip1-bound peaks were within promoters (within 2kb of transcription start sites), 7.4% were located in exons, 23.3% in introns, 0.7% in transcription termination sites, 9.7% in 5′ distal (2-50kb from a gene) regions, 3.4% in 3′ distal (2-50kb from a gene) regions, and 5.5% in intergenic (beyond 50kb from a gene) regions (Supp Fig 6d). Only 18.6% of Snip1-bound peaks were located distal to a gene (Supp Fig 6d).

**Fig. 3.**
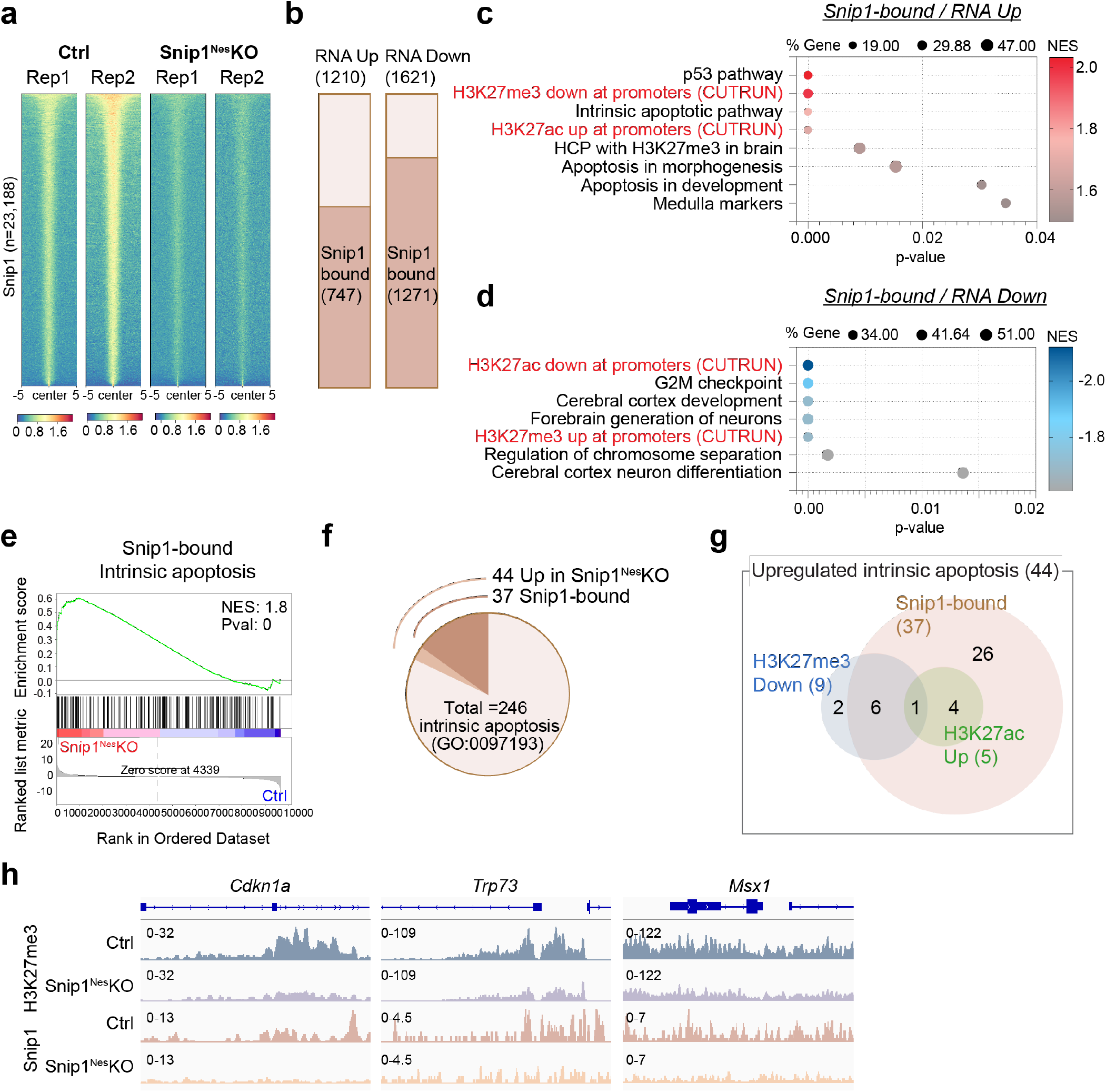
Snip1 binds to chromatin and affects H3K27me3 levels for regulating gene expression. **a** Heatmaps representing the binding intensity of two biological replicates of Snip1 CUT&RUN in control and *Snip1*^*Nes*^*-*KO NPCs. Binding intensity for 5 kb on either side of all 23,188 Snip1 CUT&RUN peaks are shown. Blue indicates low intensity and red indicates high intensity. **b** Bar charts displaying the numbers of upregulated and downregulated genes (*Snip1*^*Nes*^*-*KO vs. control, FDR <0.05) that are bound by Snip1 at their gene body. **c-d** Bubble plots of the enriched gene sets in Snip1-bound genes that became (**c**) upregulated genes and (**d**) downregulated genes in *Snip1*^*Nes*^-KO vs. control NPCs. When adding our H3K27me3/ac CUT&RUN data to the GSEA gene sets, H3K27me3 and H3K27ac levels showed anti-correlation and correlation with gene expression, respectively. **e** Representative GSEA of upregulated genes in *Snip1*^*Nes*^-KO NPCs vs. control NPCs that are bound by Snip1 in control NPCs. Intrinsic apoptosis genes were mostly Snip1-bound and were enriched in the upregulated genes in *Snip1*^*Nes*^-KO NPCs. Differentially expressed genes were first ranked by their fold-change and p-value before GSEA was performed. **f** Pie chart showing the proportions of intrinsic apoptosis genes that are upregulated in *Snip1*^*Nes*^-KO NPCs and/or bound by Snip1. **g** Venn diagram displaying the numbers of upregulated intrinsic apoptosis genes in three categories. Using our Snip1, H3K27me3, and H3K27ac CUT&RUN data, the 44 genes were categorized into 1) Snip1-bound in control NPCs, 2) reduced H3K27me3 levels in *Snip1*^*Nes*^-KO vs. control NPCs (p <0.05), and/or 3) increased H3K27ac levels in *Snip1*^*Nes*^-KO vs. control NPCs (p <0.05). **h** H3K27me3 and Snip1 CUT&RUN tracks visualized by Integrative Genomics Viewer (IGV) at upregulated intrinsic apoptosis genes. *Cdkn1a*, Chr17: 29,090,888 - 29,095,850. *Trp73*, Chr4: 154,132,565 - 154,143,373. *Msx1*, Chr5: 37,818,429 - 37,828,924.

Next, we examined whether Snip1 occupancy corelates with gene expression. Of the 1,210 upregulated genes and 1621 downregulated genes in *Snip1*^*Nes*^-KO, 747 (62%) and 1,271 (78%) were Snip1 targets, respectively, in control NPCs (Fig 3b). These overlaps are significant, with *p* = 5.55e-25 and 2.24e-155 respectively, by the hypergeometric test (given a total of 21,636 expressed genes), suggesting that Snip1 occupies these genes to regulate their expression. GSEA showed that Snip1 targets that became upregulated in *Snip1*^*Nes*^*-*KO were enriched in p53 pathway, medulla, and apoptosis (Fig 3c), whereas Snip1 targets that became downregulated in *Snip1*^*Nes*^*-*KO were enriched in G2/M checkpoint, cortical development, and chromosome segregation (Fig 3d). Intrinsic apoptosis genes were enriched in Snip1 targets that became upregulated in *Snip1*^*Nes*^-KO (Fig 3e-f, Supp Fig 6e-f). Of the 44 genes in the intrinsic apoptosis gene set, 37 gene promoters were bound by Snip1 in control NPCs (*p* = 4.62e-5; Fig 3g), suggesting that Snip1 directly suppresses these genes. These data suggest that Snip1 directly regulates genetic programs crucial to apoptosis control, cell cycle, and cortical development.

### Snip1 regulates genetic programs through H3K27 modifications

Upregulated genes in *Snip1*^*Nes*^-KO were enriched in genes whose high-CpG-density promoters that are 1) H3K27me3-occupied in the embryonic murine brain ^42^ or 2) bivalent in mouse NPCs ^46^ (Supp Fig 7a-b). This prompted us to scrutinize whether Snip1 controls genetic programs through H3K27 modifications. We profiled H3K27me3 and H3K27ac by CUT&RUN in *Snip1*^*Nes*^-KO and control Sox2-positive NPCs (Supp Fig 7c-g). We observed a strong correlation between upregulated genes, lower H3K27me3 occupancy, and higher H3K27ac occupancy, whereas downregulated genes had higher H3K27me3 occupancy and lower H3K27ac occupancy in *Snip1*^*Nes*^-KO (Fig 3c-d, Supp Fig 7h-i). Among the 44 upregulated intrinsic apoptosis genes, 9 genes showed reduced H3K27me3 levels, and 5 genes showed increased in H3K27ac levels (p <0.05, Fig 3g-h, Supp Fig 7j-k). These data suggest that Snip1 regulates genetic programs through H3K27 modifications.

### Snip1 and PRC2 co-occupy chromatin targets in NPCs

We investigated the interactions between Snip1, H3K27 methyltransferase PRC2, and histone acetyletransferases p300 and Cbp. Anti-Snip1 antibody co-immunoprecipitated with known PRC2 subunits Jarid2, Suz12, and Ezh2, but not the negative control Rbbp5 in the NPC nuclear extract (Fig 4a). Anti-Jarid2 or Ezh2 antibody co-immunoprecipitated Snip1 and other PRC2 subunits but not the negative control Rbbp5 in the NPC nuclear extract (Fig 4b-c). Anti-p300 or Cbp antibody failed to co-immunoprecipitate Snip1, suggesting that in NPCs, their physical interaction is undetectable (Supp Fig 8). Next, we performed CUT&RUN to profile Suz12 and Ezh2 in *Snip1*^*Nes*^-KO and control NPCs (Supp Fig 9a-c). Consistent with the co-IP results, Suz12, and Ezh2 co-occupy Snip1-bound target sites genome-wide (Fig 4d). The levels of Suz12, Ezh2, and H3K27me3 were significantly reduced in *Snip1*^*Nes*^-KO (Fig 4e, Supp Fig 9d-e). In contrast, H3K27ac levels were less altered in *Snip1*^*Nes*^-KO NPCs (Fig 4e, Supp Fig 9d-e). These data suggest that Snip1 binds to PRC2 to positively regulate its chromatin occupancy and H3K27me3 deposition.

**Fig. 4.**
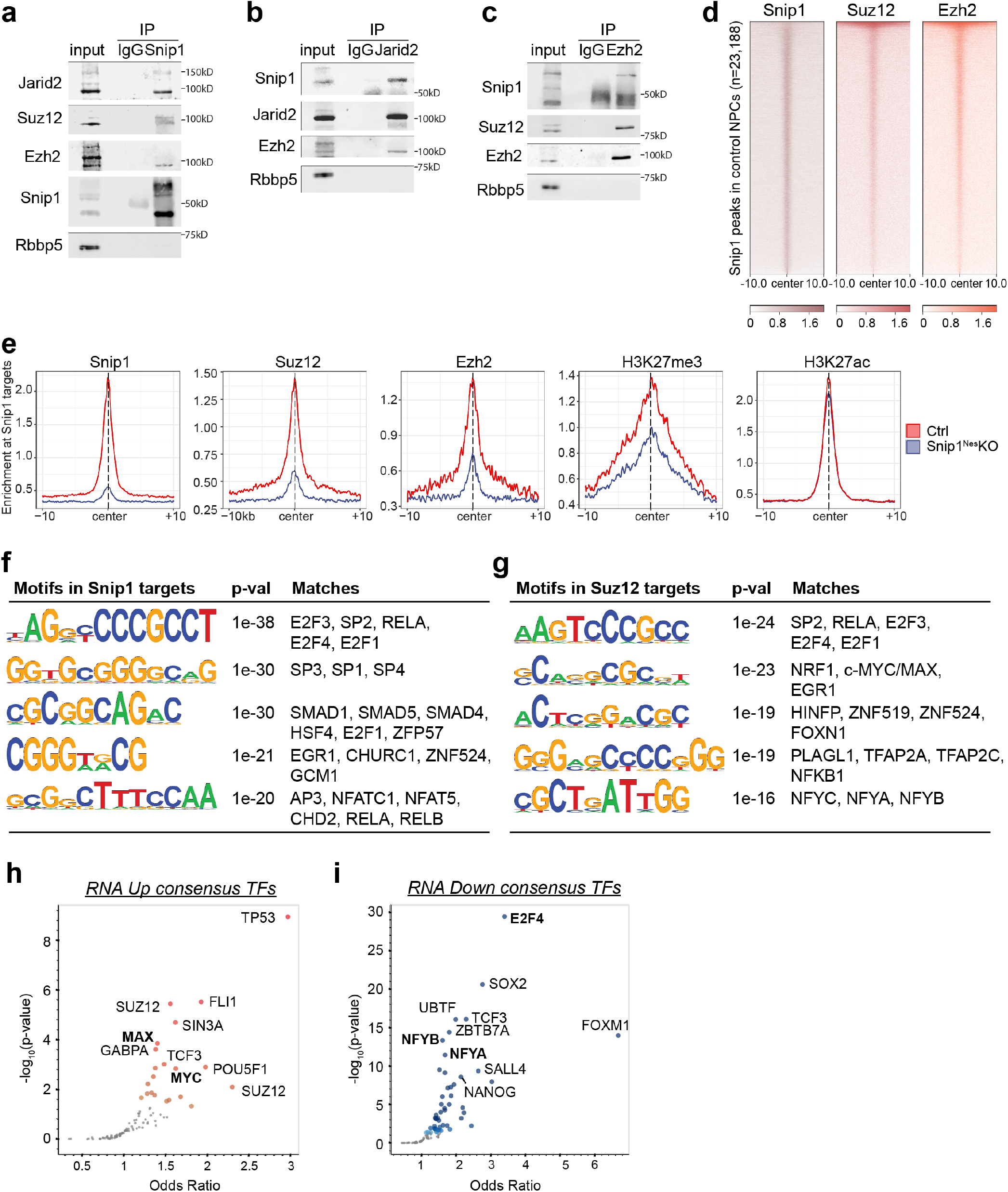
Snip1-bound regions are co-occupied by PRC2 on NPC chromatin. **a-c** Co-immunoprecipitation followed by WB to examine the interaction between Snip1 and PRC2. (**a**) Snip1, (**b**) Jarid2, or (**c**) Ezh2 was immunoprecipitated in the NPC nuclear extract. Rbbp5 was a negative control. **d** Heatmaps aligning the peaks enriched of Snip1, Suz12, and Ezh2 in control NPCs. Peak intensity for 10 kb on either side of 23,188 Snip1-bound peaks are shown. A dark color indicates high intensity and a light color indicates low intensity. **e** Profile plots comparing the median binding intensity of Snip1, PRC2, and H3K27me3/ac in *Snip1*^*Nes*^-KO vs. control NPCs at the Snip1 targets. Regions were considered true Snip1 targets when Snip1 levels reduced in *Snip1*^*Nes*^-KO vs. control NPCs with p<0.05. **f-g** Motifs of (**f**) Snip1- and (**g**) Suz12-bound regions where their levels significantly decreased in *Snip1*^*Nes*^*-*KO NPCs with fold-change >2 and p <0.05. HOMER *de novo* analysis was performed and the best five motifs which had vertebrate motif matches are listed here. **h-i** Volcano plots of transcription factors whose binding to our differentially expressed genes (FDR <0.05) has been reported. Genes were searched against ENCODE and ChEA consensus TFs from ChIP-X database using Enrichr ^48^. Darker colors show smaller p-values and large points passed p-value <0.05. Transcription factors in bold were found in our CUT&RUN motif analyses in **Fig 4f-g**.

To characterize the Snip1-PRC2 targets, we performed *de novo* motif discovery by the HOMER software ^47^. Using the criteria of fold-change >2 and p <0.05 in control vs *Snip1*^*Nes*^-KO, Snip1 targets were enriched in motifs of E2F proteins, SP1/3/4, and EGR1 (Fig 4f). Motifs of previously reported Snip1 interactors, SMAD proteins and RELA, were also found amongst Snip1 targets ^17,22^. In Suz12-bound peaks that had reduced binding in *Snip1*^*Nes*^-KO, motifs were enriched with SP2, RELA, E2F proteins, EGR1, HINFP, PLAGL1, and NF-Y subunits (Fig 4g). The similarities of Snip1- and Suz12-bound motifs point to potential interactions of Snip1 and PRC2 with some of these transcription factors.

We next examined whether the Snip1- or Suz12-bound motifs were overrepresented in upregulated or downregulated genes. Using Enrichr ^48^, we found that upregulated genes in *Snip1*^*Nes*^-KO were targets of TP53, FLI1, SUZ12, MAX, and MYC, whereas downregulated genes were targets of E2F4, SOX2, NFYB and NFYA (Fig 4h-i) E2F proteins were uncovered by motif and Enrichr analyses, and E2F4 targets *Mcm7* and *Anp32e* were Snip1 and PRC2 targets that had reduced binding in *Snip1*^*Nes*^-KO (Fig 4i, Supp Fig 9f). Among upregulated genes in *Snip1*^*Nes*^-KO, MYC and MAX targets were identified by both motif and Enrichr analyses (Fig 4h, Supp Fig 9g). These data suggest that E2F proteins, MYC, MAX, and Snip1-PRC2 may co-regulate genes.

### Snip1 functionally interacts with PRC2 to modulate the intrinsic apoptosis program

We tested whether Snip1 functions cooperatively or antagonistically with PRC2 to regulate NPC survival *in vivo*. To genetically deplete the PRC2 core subunit Eed, we used *Nes*:Cre to excise exons 3 to 6 of *Eed* in *Eed*^*Nes*^-KO and *Snip1*^*Nes*^*-Eed*^*Nes*^-dKO (Supp Fig 10a-b). At E13.5, the Eed depletion alone was not sufficient to induce apoptosis or alter self-renewal of NPCs (Supp Fig 10c-e). We examined apoptosis and neurogenesis in *Snip1*^*Nes*^*-Eed*^*Nes*^-dKO brains at E13.5 (Fig 5a). Compared with *Snip1*^*Nes*^*-* KO, *Snip1*^*Nes*^*-Eed*^*Nes*^-dKO had significantly fewer cl-caspase 3-positive cells (Fig 5b-c), more Sox2-positive NPCs (Fig 5d-e), and more Tbr2- or Insm1-positive intermediate progenitors (Fig 5f-i). Tuj1-positive immature neurons were not markedly affected by the Snip1–PRC2 functional interaction (Supp Fig 10f-g). Eed depletion in the *Snip1*^*Nes*^*-*KO embryonic brain reduced apoptosis and rescued NPCs and intermediate progenitors. Therefore, Snip1 likely counterbalances PRC2 in regulating intrinsic apoptosis in the developing brain.

**Fig. 5.**
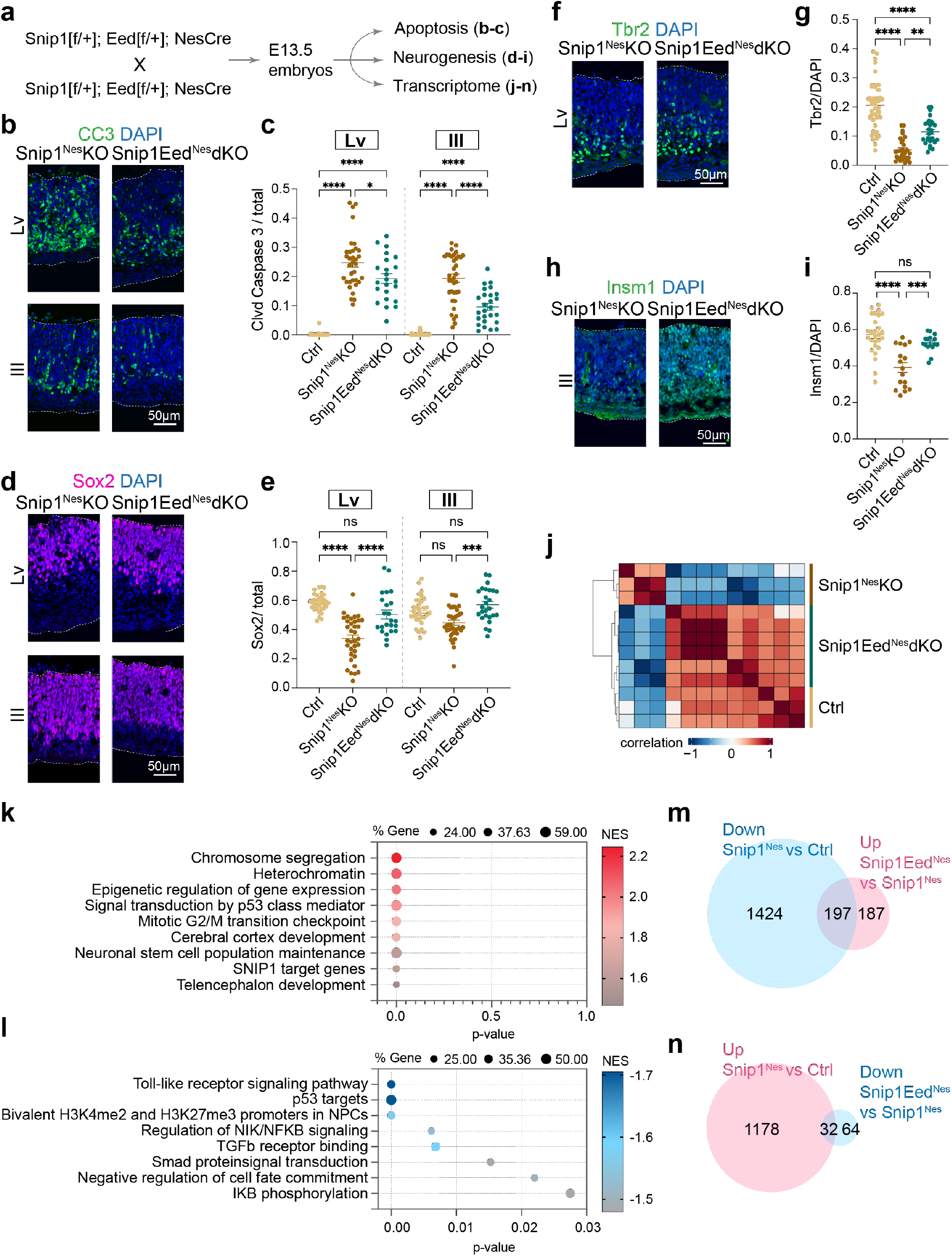
Eed depletion reduces apoptosis and brain dysplasia of the *Snip1*^*Nes*^-KO brain. **a** Schematic of the genetic cross depleting both *Snip1* and *Eed* and its downstream assays. **b-c** IF analysis of CC3 overlayed with DAPI of the E13.5 brain. Bar, 50 μm. Each data point represents one image. N=8 embryos for control, n=7 for *Snip1*^*Nes*^-KO, and n=5 for *Snip1*^*Nes*^*-Eed*^*Nes*^-dKO. **d-k** IF of NPC marker (**d**) Sox2 and neural lineage markers (**f**) Tbr2, (**h**) Insm1, or (**j**) Tuj1 overlayed with DAPI of the E13.5 brain. Bar, 50 μm. The populations of (**e**) Sox2-positive, (**g**) Tbr2-positive, or (**i**) Insm1-positive cells in the neuroepithelial lining of lateral and/or 3^rd^ ventricles were quantified. Each data point represents one image. N=5-8 embryos for control, n=3-7 for *Snip1*^*Nes*^-KO, n=3-5 for *Snip1*^*Nes*^*-Eed*^*Nes*^-dKO. For Panels **c, e, g**, and **i**, data are presented as mean ± SEM, and two-way ANOVA was used for statistical analysis. ns = not statistically significant; *p <0.05; **p <0.01; ***p <0.001; ****p <0.0001. **j** Unsupervised clustering of RNA-seq data from control (n=3), *Snip1*^*Nes*^-KO (n=3), and *Snip1*^*Nes*^*-Eed*^*Nes*^-dKO (n=6) brains at E13.5. RNAs from forebrain and midbrain regions were sequenced and merged for the downstream analyses. Blue indicates a negative correlation and red indicates a positive correlation. **k-l** Bubble plots of the enriched gene sets in (**k**) upregulated genes and (**l**) downregulated genes in *Snip1*^*Nes*^*-Eed*^*Nes*^-dKO vs. *Snip1*^*Nes*^-KO brain cells. Differentially expressed genes were first ranked by their fold-change and p-value before GSEA was performed. **m-n** Venn diagrams displaying the numbers of differentially expressed genes with FDR <0.05. The lists of downregulated genes in *Snip1*^*Nes*^-KO vs. control and upregulated genes in *Snip1*^*Nes*^*-Eed*^*Nes*^-dKO vs. *Snip1*^*Nes*^-KO are compared in (**m**). The lists of upregulated genes in *Snip1*^*Nes*^-KO vs. control and downregulated genes in *Snip1*^*Nes*^*-Eed*^*Nes*^-dKO vs. *Snip1*^*Nes*^-KO are compared in (**n**).

We profiled the transcriptomes of control, *Snip1*^*Nes*^*-*KO, and *Snip1*^*Nes*^*-Eed*^*Nes*^-dKO brain tissues. Unsupervised clustering based on top 3000 most variant genes (based on median variation values) suggest a higher similarity in the transcriptomes between control and *Snip1*^*Nes*^*-Eed*^*Nes*^-dKO compared with *Snip1*^*Nes*^*-*KO (Fig 5j). Using fold-change >2 and p <0.05 to compare datasets from *Snip1*^*Nes*^*-KO* and *Snip1*^*Nes*^*-Eed*^*Nes*^-dKO, we identified 184 upregulated genes and 994 downregulated genes in *Snip1*^*Nes*^*-Eed*^*Nes*^-dKO (Supp Fig 10h). Upregulated genes in *Snip1*^*Nes*^*-Eed*^*Nes*^-dKO were enriched in functions related to chromosome segregation, G2/M transition checkpoint, cerebral cortex/telencephalon development, and heterochromatin (Fig 5k). Downregulated genes in *Snip1*^*Nes*^*-Eed*^*Nes*^-dKO were enriched in functions related to negative regulation of cell fate commitment, bivalent promoters in NPCs, p53 targets, and TLR, NFκB, and TGFβ signaling pathways (Fig 5l). Interestingly, upregulated genes in *Snip1*^*Nes*^*-Eed*^*Nes*^-dKO (versus *Snip1*^*Nes*^-KO) were enriched in functions related to chromatin modifications and remodeling (Supp Fig 10i), suggesting that the Snip1-PRC2 interaction regulates chromatin organization.

To identify a potential molecular explanation of the rescue of NPCs and intermediate progenitors in *Snip1*^*Nes*^*-Eed*^*Nes*^-dKO, we analyzed differentially expressed genes among control, *Snip1*^*Nes*^*-Eed*^*Nes*^-dKO, and *Snip1*^*Nes*^*-KO*. One hundred and ninety-seven downregulated genes in *Snip1*^*Nes*^*-*KO partially regained expression in *Snip1*^*Nes*^*-Eed*^*Nes*^-dKO (Fig 5m). These rescued genes are involved in G2/M checkpoint, E2F targets, cholesterol homeostasis, mTORC1 signaling, androgen response, Myc targets, fatty acid metabolism, and Wnt signaling (Supp Fig 10j). In contrast, 32 upregulated genes in *Snip1*^*Nes*^*-KO* became downregulated in *Snip1*^*Nes*^*-Eed*^*Nes*^*-KO* (Fig 5n) and are involved in p53 pathway, inflammatory and interferon gamma responses, NFκB signaling, and apoptosis (Supp Fig 10k). These data suggest that PRC2 and Snip1 interact to regulate apoptosis, cell cycle progression, neuronal differentiation, and chromatin organization to maintain the appropriate balance of NPC division, apoptosis, and differentiation in the developing brain.

## DISCUSSION

In the developing brain, neural cells are prone to undergo apoptosis ^1-4^. Using embryonic mouse brains as a model, we uncovered that Snip1 promotes neurogenesis and cortical development and suppresses intrinsic apoptosis. Snip1 and PRC2 co-occupy chromatin targets to co-regulate select genetic programs. Initially, we had hypothesized that Snip1 directly suppresses apoptotic genes in a cooperative manner with PRC2. However, the rescue of *Snip1*^*Nes*^-KO NPCs by Eed/PRC2 depletion came as an intriguing surprise and suggest that Snip1 and PRC2 have a balancing relationship for regulating genetic programs in the brain.

A role of Snip1 in apoptosis has been implicated by other studies. Snip1 depletion induced acridine orange (apoptosis detection) signals in zebrafish embryos ^31^. SNIP1 depletion in a human osteosarcoma cell line U2OS triggers γH2AX increases ^49^. Our study advances our knowledge by uncovering (1) the participation of caspases 9 and 3, (2) the chromatin-based role of Snip1, (3) the delicate balancing relationship between Snip1 and PRC2 to fine-tune genetic programs, (4) implicating the participation of E2F proteins, SMADs, and RELA/B, and (5) the requirement of Snip1 for PRC2 binding to chromatin and H3K27me3 deposition in NPCs. Nevertheless, the precise mechanisms by which Snip1–PRC2 localizes to chromatin targets and how the downstream factors orchestrate caspase 9-dependent apoptosis remain unclear and subject to extensive future studies.

The role of caspase 9 and intrinsic apoptosis in brain development has been little understood, despite the human genetic evidence connecting loss of function in caspase 9 to neural tube defects ^50-52^ and pediatric brain tumors ^53,54^. Global KO of either caspase 3 or 9 in mice leads to prenatal death with brain malformation, including neural tube closure defect and exencephaly ^55-57^. These findings point to a conserved requirement of caspase 3 and 9 for brain development. Our study implicates the essential roles of Snip1 and PRC2 in dampening the activities of caspase 9 and caspase 3 in order to balance the division and death of NPCs. And this dampening is potentially mediated through p53, whose protein levels markedly increase and coincide with higher expression of p53 targets in *Snip1*^*Nes*^-KO NPCs.

Our study identifies a physiological role of Snip1 in regulating cell cycle that cell line studies have previously proposed ^27,58,59^. We showed that *Snip1*^*Nes*^-KO NPCs have defective genetic programs in chromosome segregation, cell proliferation, and replication fork. In culture, *Snip1*^*Nes*^-KO NPCs diminished the growth of neurospheres over time. We previously showed that PRC2 maintains the self-renewal of NPCs by suppressing neurogenesis ^60^. This study uncovered that Snip1 and PRC2 transcriptionally regulate cell cycle progression; this may be clinically relevant to developmental defects including skull dysplasia, global developmental delay, and intellectual disability and seizure that are associated with 1097A>G (Glu366Gly) variant of *SNIP1* ^61,62^. Our findings about the Snip1–PRC2 interactions in NPC survival and functions improves our understanding about the neurodevelopmental disorders caused by *SNIP1* mutations and PRC2 dysfunction.

## ACKNOWLEDGEMENTS

The authors thank V. Shanker for editing the manuscript; J. Houston and K. Lowe for FACS; M. Evans, I. Lam, and I. Chapman for experimental assistance. RNAscope was performed by the Comparative Histology Core at SJCRH. Sequencing was performed at the Harwell Center for Biotechnology and images were acquired at the Cell & Tissue Imaging Center, both of which are supported by SJCRH and NCI P30 (CA021765). M.N.D, Y.F., and B.X. are supported by NCI P30 grant (CA21765). This work is funded by the American Lebanese Syrian Associated Charities, American Cancer Society (132096-RSG-18-032-01-DDC), and NIH (1R01GM134358-01). The content is solely the responsibility of the authors and does not necessarily represent the official views of the National Institutes of Health. The funders had no role in study design, data collection and analysis, decision to publish, or preparation of the manuscript.

## AUTHOR CONTRIBUTIONS

Y.M.: Most experiments, data analyses, and manuscript writing. M.N.D. and B.X.: bioinformatics analyses. K.L.: mouse harvesting and rescue by lentiviral Snip1. P.S.: ideas on studying apoptosis and inhibitor assays. N.C. and H.C.: mouse breeding and DNA constructs. Y.F.: supervision of bioinformatics analyses. J.C.P.: project design, data analyses, and manuscript writing with inputs from all authors.

## METHODS

### Buffers

PBS: 137 mM NaCl, 2.7 mM KCl, 10 mM phosphate buffer (pH 7.4)

PBST: PBS with 0.1% Triton X-100

HEPM (pH 6.9): 25 mM HEPES, 10 mM EGTA, 60 mM PIPES, 2 mM MgCl_2_

IF blocking buffer: 1/3 Blocker Casein (ThermoFisher 37528), 2/3 HEPM with 0.05% Triton X-100

Buffer A: 10 mM HEPES (pH 7.9), 10 mM KCl, 1.5 mM MgCl_2_, 0.34 M sucrose, 10% glycerol

Buffer D: 400 mM KCl, 20 mM HEPES, 0.4 mM EDTA, 20% glycerol iDISCO PTx.2: PBS with 0.2% Triton X-100

iDISCO PTwH: PBS with 0.2% Tween-20 and 10 μg/mL heparin

iDISCO Permeabilization solution: PTx.2 with 306 mM glycine and 20% DMSO

iDISCO Blocking solution: PTx.2 with 6% donkey serum and 10% DMSO

CUT&RUN Binding buffer: 20 mM HEPES-KOH (pH 7.9), 10 mM KCl, 1 mM CaCl_2_, 1 mM MnCl_2_

CUT&RUN Wash buffer: 20 mM HEPES (pH 7.5), 150 mM NaCl, 0.5 mM spermidine, Protease Inhibitor Cocktail (Sigma-Aldrich 11873580001)

CUT&RUN Digitonin block buffer: CUT&RUN Wash buffer with 2 mM EDTA and 0.05% digitonin CUT&RUN 2X Stop buffer stock: 340 mM NaCl, 20 mM EDTA, 4 mM EGTA, 0.02% digitonin CUT&RUN Stop buffer: Into 1 mL of 2X Stop buffer stock, add 5 μL of 10 mg/mL RNase A and 133 μL of 15 mg/mL GlycoBlue™ Coprecipitant (ThermoFisher AM9516)

Antibodies used in this study are listed in Supplementary Table 1.

### Animals

All animal experiments were approved by the Institutional Animal Care and Use Committee at St. Jude Children’s Research Hospital and were conducted in accordance with ethical guidelines for animal research. To generate conditional knockout embryos, we used the Cre/lox system. *Snip1*-tm1a (Infrafrontier/EMMA 04224) were first crossed with *Actin*-FLPe to generate the *Snip1*-flox line. To genetically label NPCs, *Snip1*-flox; *Nestin*-Cre mice were crossed with *Sox2*-eGFP transgenic mice. Embryos were harvested at embryonic days as indicated in the figures or text. For all the animal experiments, both sexes were included. The following mouse lines were used in this study and genotyping primers and conditions are shown in Supplementary Table 2:

*Snip1*-tm1a: B6Dnk;B6N-*Snip1*<tm1a(EUCOMM)Wtsi>/H (Infrafrontier/EMMA 04224)

*Eed*-flox: B6;129S1-*Eed*^tm1Sho^/J (JAX Stock 022727) ^63^

*Actin*-FLPe: B6;SJL-Tg(*ACT*FLPe)9205Dym/J (a gift from Dr. Peter McKinnon at St. Jude Children’s

Research Hospital, JAX Stock 003800) ^64^

*Nestin*-Cre: B6.Cg-Tg(*Nes*-Cre)1Kln/J (JAX Stock 003771) ^65^

*Emx1*-Cre: B6.129S2-Emx1^tm1(cre)Krj^/J (a gift from Dr. Peter McKinnon at St. Jude Children’s Research

Hospital, JAX Stock 005628) ^66^

*Sox2*-eGFP: B6;129S1-*Sox2*^tm1Hoch^/J (JAX Stock 017592) ^67^

### Isolation and Culturing of Mouse NPCs

To obtain mouse brain cells, embryos at an indicated embryonic day were dissected out from the uterus and visceral yolk sac. A part of the tail or limbs was collected for genotyping. Brains were dissected from embryos under the dissection microscope in cold 1x PBS. Then, 300 μL Dulbecco’s Modified Eagle’s Medium (DMEM) (ATCC 30-2002) and 150 μL of 10 mg/mL collagenase Type II (Worthington LS004176) were added to each brain and incubated for 5-10 min at 37 °C. After centrifugation at 1000 xg for 3 min, the tissue was incubated with 500 μL 0.25% Trypsin-EDTA (ThermoFisher 25200056) for 5 min at 37 °C. Trypsinization was quenched with 500 μL DMEM supplemented with 10% fetal bovine serum (FBS) and pelleted by centrifugation at 1000 xg for 3 min. Alternatively, cells were dissociated from the brain using the papain dissociation system (Worthington LK003153). Cells were then resuspended in 500 μL of NPC culture media (NeuroCult™ Proliferation Media; STEMCELL Technologies 05702) supplemented with 30 ng/mL human recombinant epidermal growth factor (rhEGF) (STEMCELL Technologies 78006) and filtered through a 40 μm filter (Fisherbrand™ 22-363-547) to obtain single cells. To collect NPCs, the dissociated brain cells were cultured in ultra-low attachment 6-well plates (Corning^®^ Costar^®^ CLS3471) in the NPC culture media at 37 °C. NPCs formed neurospheres in suspension. For passaging, neurospheres were incubated with Accutase™ (STEMCELL Technologies 07920) for 5 min at 37 °C and then dissociated by pipetting. After adding an equal volume of the NPC culture media, dissociated cells were centrifuged at 500 xg for 5 min. Cells were then grown in the NPC culture media either in suspension in the ultra-low attachment 6-well plates or on matrigel (Corning™ 354230) coated plates. The medium was changed every 2 to 3 days. For collecting uncultured NPCs, cells from the Sox2-eGFP brains or stained with NeuroFluor™ CDr3 (STEMCELL Technologies 01800) were sorted by fluorescence-activated cell sorting (FACS).

### Neurosphere Assay

NPCs were seeded into ultra-low attachment 6-well plates and grown in the NPC culture media at 37 °C with 250 μL of media added every 2 days. Neurospheres were imaged after 5 days of culturing. For a neurosphere rescue experiment, freshly dissociated brain cells were first transduced with lentivirus delivering human SNIP1 transgene (System Biosciences CD823A-1) in ultra-low attachment 6-well plates. After 5 days, transduced neurospheres were then dissociated and seeded for a neurosphere assay. All the neurosphere assays were done in three replicates. At least 8 images of neurospheres per well were captured with ZEISS AxioObserver D1 at 5x magnification. The area and the number of neurospheres were quantified by using FIJI. To generate clean binary images, images were processed with “Process” -> “Find edges” followed by “Image” -> “Adjust” -> “Threshold”. After inverting the images, the number and the area of the neurospheres were obtained by selecting “Analyze” -> “Analyze particles”. If multiple neurospheres were too close for the software to quantify individually, one of the two methods was applied after generating the binary images: manual quantification by drawing the outline of each neurosphere and selecting “Analyze” -> “Measure”, or computationally separating the neurospheres by selecting “Process” -> “Binary” -> “Fill Holes,” followed by “Process” -> “Binary” -> “Watershed”.

### Inhibitor Treatment and FACS-Based Cell Death Assay

5 × 10^5^ NPCs from Snip1[+/+] and Snip1[flox/flox] embryos were seeded onto each well of matrigel-coated 6-well plates. On the following day, cells were incubated with mCherry-Cre lentivirus (Vector Core Lab at St. Jude Children’s Research Hospital) for 8 hours, washed twice with 1X PBS, and cultured for 3 days. To quantify the population of cells with active caspases 3 and 7, cells were incubated at 37 °C with reconstituted FAM-FLICA^®^ at a 1:300 dilution (ImmunoChemistry Technologies 94) for 30 min. Cells were fixed in a 4% formaldehyde solution at room temperature for 15 min and washed twice with 1X PBS. FAM-FLICA–positive cells were quantified by FACS (Excitation: 492 nm, Emission: 520 nm). FACS data were analyzed by FlowJo. To examine whether cell death is via activation of caspase 8 or 9, Z-IETD-FMK (a caspase 8 inhibitor) and Z-LEHD-FMK TFA (a caspase 9 inhibitor) were dissolved in DMSO at 50mM (Compound Management Center at St. Jude Children’s Research Hospital). After cells were incubated with mCherry-Cre lentivirus for 8 hours, these compounds were added at a series of concentrations and incubated for 3 days before FACS analysis. For all the inhibitor treatment assays, medium with the inhibitors was changed every 2 days.

### Subcellular Protein Extraction

After washing cells once with 1x PBS, they were resuspended in 2x volume of Buffer A supplemented with PI, 1 mM DTT, and 0.1% TritonX-100 and placed on ice for 5 min. Cells were centrifuged at 1750 x g for for 2 min at 4 °C and supernatant was collected as the cytoplasmic fraction. The nuclear pellet was then resuspended in 1x volume of Buffer D supplemented with PI, 1 mM DTT, and 0.1% TritonX-100 and placed on ice for 30 min (If volumes were large, tubes were rotated at 4 °C). The lysate was centrifuged at 1750 x g for for 2 min at 4 °C and supernatant was collected and diluted with an equal volume of H_2_O (nuclear fraction). The chromatin pellet was washed once with cold 1x PBS and resuspended in 1x volume of 0.1 N HCl at 4 °C overnight (O/N). The supernatant was neutralized with the equal volume of 1.5 M Tris-HCl (pH 8.8) (chromatin fraction). For whole-cell lysate extraction, cells were washed once with 1X PBS and resuspended directly in 2x volume of Buffer D supplemented with PI, 1 mM DTT, and 0.1% TritonX-100.

### Co-Immunoprecipitation

Nuclear proteins were extracted as above. Then, 15 μL of protein A and 15 μL of protein G Dynabeads™ (Invitrogen 10002D and 10004D) were washed once with 1x PBST. For pre-bound co-immunoprecipitation, the beads were resuspended in 100 μL of HEPM and 4 μg primary antibody was added. The tube was gently shaken at room temperature for 2 h. The beads were washed once with 1x PBST, and approximately 2.5 mg of nuclear extract was added to the antibody-prebound beads and the tube was rotated at 4 °C for 2.5-4 hours. Beads were then washed three times with 1x PBST and proteins were eluted with 0.1 M glycine (pH 2.3) at room temperature. Eluates were neutralized with 1/10 volume of 1.5 M Tris-HCl (pH 8.8). For co-immunoprecipitation with free antibodies, approximately 2.5 mg of nuclear extract was incubated with 4 μg of primary antibody and rotated at 4 °C for 4 hours. Beads were then added to the extract and gently shaken for 1 hour at room temperature. The same washing and elution steps were performed as for the pre-bound co-immunoprecipitation.

### Western Blotting (WB)

For SDS-PAGE, resolving and stacking gels were prepared using the following composition. Resolving gels: 6-12% ProtoGel (National Diagnostics EC8901LTR), 0.375 M Tris-HCl (pH 8.8), 0.1% SDS, 0.1% ammonium persulfate (APS) and 0.1% TEMED (National Diagnostics EC-503). Stacking gels: 3.9% ProtoGel, 0.125M Tris-HCl (pH 6.8), 0.1% SDS, 0.05% APS, and 0.12% TEMED. After proteins were separated by SDS-PAGE, they were transferred onto a 0.45 μm nitrocellulose membrane (Bio-Rad 1620115) by the semi-dry transfer system. Membranes were blocked with 2% BSA in HEPM for 1hour at room temperature and incubated in primary antibodies diluted in the 2% BSA at 4 °C O/N. On the following day, the membrane was washed three times with 1x PBST and incubated in IRDye®-conjugated secondary antibodies (LI-COR) or Clean-Blot™ IP detection reagent (ThermoFisher 21230) on a shaker for 1 hour at room temperature. The membrane was washed three times with 1x PBST and immediately imaged on an Odyssey® Fc imaging system (LI-COR). The membrane stained with Clean-Blot™ IP detection reagent was treated with SuperSignal™ West Pico PLUS Chemiluminescent Substrate (ThermoFisher 34577) for at least 5 min at room temperature before imaging. Signals were quantitated using the Image Studio™ software (version 1.0.14; LI-COR).

### BrdU Administration

Mice were administered 5-bromo-2′-deoxyuridine (BrdU, Sigma-Aldrich B5002) reconstituted in sterile 1x PBS by intraperitoneal injection at a dose of 50 mg/kg. After 5 hours, the mice were sacrificed for dissection.

### Cryosection

Mouse embryos at an indicated embryonic day were fixed in 4% formaldehyde at 4 °C O/N. The embryos were washed three times in 1x PBS for 30 min at room temperature and placed in 15% sucrose diluted in 1x PBS at room temperature until the embryos sank to the bottom of the tube. Then, the embryos were moved to a 30% sucrose solution and incubated at room temperature until the embryos sank. The embryos were then treated in the embedding medium Tissue-Tek® O.C.T. Compound (Sakura Finetek USA INC 4583) to rinse the residual sucrose. Each embryo was mounted in the embedding media in a cryosection mold placed on dry ice and 12-μm sagittal cryosections were obtained (Leica CM3050 S).

### Immunofluorescence (IF)

For cryosections, the slides were permeabilized in 1x PBST at 4 °C O/N. After drawing the outline of the staining area with a hydrophobic barrier pen (ImmEdge® H-4000), the slides were blocked in the IF blocking buffer for 2-3 hours at room temperature. Primary antibodies were diluted at the optimized concentration in the IF blocking buffer and incubated at 4 °C O/N. Sections were washed three times with 1x PBST and incubated with the fluorescent dye–conjugated Alexa Fluor secondary antibodies (Invitrogen) diluted at 1:500 for 2-3 hours at room temperature. Sections were washed three times with 1x PBST and incubated in 1 mg/mL DAPI (Sigma-Aldrich D9542) diluted at 1:500 in 1x PBS at room temperature for 1 hour. Finally, the sections were washed once with 1x PBS and mounted with ProLong™ Gold Antifade Mountant (Life Technologies P10144). For detecting BrdU, the tissue was permeabilized as above and rinsed with 1x PBS for 5 min at room temperature. Then, the tissue was treated with 2 N HCl for 1 hour at room temperature. Sections were washed multiple times with 1x PBS to remove all traces of HCl and were blocked and stained as above. Images were acquired with a Nikon C2 laser scanning confocal microscope.

### Image Analysis

All of the IF image analyses were performed by FIJI. For counting the number of cells, the images were first converted to an 8-bit grayscale. In the automatic nuclei counter plugin (ITCN) of FIJI, for each primary antibody, “Width”, “Minimum Distance” and “Threshold” were set manually based on the area and the intensity of the signal on the representative images. The same parameters on ITCN were applied for control and experimental groups. For each group, at least five images were analyzed. For measuring the thickness of a tissue, distance between two hand-drawn lines on an image was measured. The publicly available InteredgeDistance macro was used for calculating the average of the shortest distances of randomly selected points on the two lines.

### iDISCO

We performed iDISCO clearing method using the protocol developed by the Tessier-Lavigne lab ^68^. In brief, mouse embryos at an indicated embryonic day were fixed in 4% formaldehyde and washed in 1X PBS (see Cryosection section). Samples were first dehydrated by incubating them for 1 hour each with the increasing concentrations of methanol at 20%, 40%, 60%, 80%, and 100% at room temperature. Samples were then incubated in 66% dichloromethane (DCM)/ 33% methanol O/N. Samples were washed twice in 100% methanol, pre-chilled at 4 °C and incubated with 5% H_2_O_2_ in methanol at 4 °C O/N. Samples were then rehydrated by incubating them for 1 hour each with the decreasing concentrations of methanol at 80%, 60%, 40%, 20% and 1X PBS at room temperature. Samples were washed twice in PTx.2 for 1 hour each. For immunolabeling, samples were incubated in a permeabilization solution at 37 °C for 2 days and blocked in Blocking solution at 37 °C for 2 days.

Primary antibodies diluted in PTwH supplemented with 5% DMSO and 3% donkey serum were incubated at 37 °C for 3-4 days. Samples were moved to the freshly diluted antibodies and incubated for another 3 - 4 days. Samples were washed in PTwH for 4-5 times for the entire day and incubated with the secondary antibodies diluted in PTwH supplemented with 3% donkey serum at 37 °C for 3-4 days. Again, the antibodies were replaced with freshly diluted antibodies and incubated for an additional 3-4 days. Samples were washed in PTwH for 4-5 times for the entire day. For clearing, samples were first dehydrated with methanol as described above and treated with 66% DCM/ 33% methanol for 3 hours at room temperature. Samples were washed twice with 100% DCM to rinse off methanol and incubated with dibenzyl ether to clear the tissues. Images were acquired with a LaVision light sheet microscope.

### RNAscope VS Duplex Assay

To detect the expression pattern of transcripts in the brain, mouse embryos at an indicated embryonic day were fixed in 10% neutral-buffered formalin (NBF) at room temperature. Fixed embryos were paraffin-embedded and sectioned at a thickness of 4 μm. RNA probes were designed and purchased from ACDBio; *Snip1* probe targeting 1270-2274 bp of NM_001356560.1 and *Eomes/Tbr2* probe targeting 1289-2370 bp of NM_010136.3 (Cat. 429649-C2). Sectioning and *in situ* hybridization (ISH) were done by the Comparative Histology Core at St. Jude Children’s Research Hospital by following manufacturer’s instructions. Brightfield images were acquired with Keyence BZ-X700.

### RNA Extraction and Reverse Transcription

Total RNA was extracted from FACS-sorted cells using TRIzol reagent (Invitrogen™ 15596026) and Direct-zol™ RNA Microprep (Zymo Research R2062) by following manufacturer’s instructions. DNA digestion with DNase I was also performed as part of the RNA extraction. cDNA was prepared with 500-1000 ng of total RNA using SuperScript™ IV VILO™ Master Mix (ThermoFisher 11766050) by following manufacturer’s instructions.

### Real-Time Quantitative PCR (RT-qPCR)

RT-qPCR was performed with PowerUp™ SYBR™ Green Master Mix (Applied Biosystems™ A25778) using Applied Biosystems QuantStudio 3. Primers are listed in Supplementary Table 3. Three technical replicates were set up for each gene target. For data analysis, 2^-ΔΔCt^ method, which compares the difference in the threshold cycle values of control and experimental samples, was used. The threshold cycle values of a gene of interest was normalized to that of the housekeeping gene *Gapdh*.

### RNA-Seq analysis

Paired-end 100-cycle sequencing was performed on NovaSeq6000 sequencer by following the manufacturer’s instructions (Illumina). Raw reads were first trimmed using TrimGalore (version 0.6.3) available at: https://www.bioinformatics.babraham.ac.uk/projects/trim_galore/, with parameters ‘-- paired --retain_unpaired’. Filtered reads were then mapped to the *Mus musculus* reference genome (GRCm38.p6 + Gencode-M22 Annotation) using STAR (version 2.7.9a) ^69^. Gene-level read quantification was done using RSEM (version 1.3.1) ^70^. To identify the differentially expressed genes between control and experimental samples, the variation in the library size between samples was first normalized by trimmed mean of *M* values (TMM) and genes with CPM < 1 in all samples were eliminated. Then, the normalized data were applied to linear modeling with the voom from the limma R package ^71^. Gene set enrichment analysis (GSEA) was performed against using the MSigDB database (version 7.1), and differentially expressed genes were ranked based on the their log_2_(FC) * -log_10_(p-value) ^72,73^.

### CUT&RUN

Approximately 3 × 10^5^ NPCs sorted for *Sox2*-eGFP were mixed with 3 × 10^4^ *Drosophila* S2 cells per reaction. We performed CUT&RUN using the protocol developed by the Henikoff lab ^43^. In brief, Bio-Mag®Plus Concanavalin-A (Con A) coated beads (Bangs Laboratories BP531) were washed and activated with Binding buffer. Nuclei of NPCs with S2 spike-in were gently prepared (see Subcellular Protein Extraction section). Activated Con A beads and nuclei were then mixed and rotated for 5 min at room temperature. They were then blocked with Digitonin block buffer for 5 min at room temperature. 0.5-1 μg of primary antibody with 0.25 μg Spike-in antibody (Active Motif 61686) diluted in Digitonin block buffer was added to the bead-nuclei mixture and incubated for 3 hours (histone marks) or O/N (the rest) at 4 °C. Beads were washed three times with Digitonin block buffer and incubated with pA-MNase for 1 hour at 4 °C. Beads were washed three times with Wash buffer and incubated in Wash buffer for 10 min on ice. The pA-MNase was activated by incubating the beads with 2 mM CaCl_2_ for 25 min on ice and quenched by adding the Stop buffer. DNA was released from the beads by incubating them for 30 min at 37 °C and collected by centrifugation at 16,000 x g for 5 min at 4 °C. DNA was then isolated by using a phenol/chloroform extraction method. Libraries were constructed using ACCEL-NGS® 1S Plus DNA Library Kit by following the manufacturer’s instructions (Swift Biosciences 10024). Purified libraries were analyzed with TapeStation (Agilent), using the High Sensitivity D1000 reagents (Agilent 5067-5585) before sequencing. IgG primary antibody was used as the negative control.

### CUT&RUN Analysis

CUT&RUN libraries were sequenced on NovaSeq6000 sequencer and generated 50 bp paired-end reads. The reads were aligned to mouse mm10 genome reference and fruit fly dm6 genome reference by BWA (version 0.7.170.7.12, default parameter). Duplicated reads were marked by the bamsormadup from the biobambam tool (version 2.0.87) available at https://www.sanger.ac.uk/tool/biobambam/. Uniquely mapped reads were kept by samtools (parameter “-q 1 -F 1804,” version 1.14). Fragments < 2000 bp were kept for peak calling and bigwig files were generated for visualization. SICER ^44^ and macs2 ^45^ were both used for peak calling, to identify both the narrow and broad peak correctly. With SICER, we assigned peaks that were at the top 1 percentile as the high-confidence peaks and the top 5 percentile as the low-confidence peaks. Two sets of peaks were generated: Strong peaks called with parameter ‘FDR < 0.05’ by at least one method (macs2 or SICER) and weak peaks called with parameter ‘FDR < 0.5’ by at least one method (macs2 or SICER). Peaks were considered reproducible if they were supported by a strong peak from all replicates or at least one strong peak and a weak peak in the other replicates. For downstream analyses, heatmaps were generated by deepTools ^74^ and gene ontology was performed with Enrichr ^48,75^ and GSEA, in addition to custom R scripts. For differential peak analysis, peaks from two replicates were merged and counted for number of overlapping extended reads for each sample (bedtools v2.24.0) ^76^. Then we detected the differential peaks by the empirical Bayes method (eBayes function from the limma R package) ^71^. For downstream analyses, heatmaps were generated by deepTools (v3.5.0) ^77^. Peaks were annotated based on Gencode following this priority: “Promoter.Up”: if they fall within TSS - 2kb, “Promoter.Down”: if they fall within TSS - 2kb, “Exonic” or “intronic”: if they fall within an exon or intron of any isoform, “TES peaks”: if they fall within TES ± 2kb, “distal5” or “distal3” if they are with 50kb upstream of TSS or 50kb downstream of TES, respectively, and they are classified as “intergenic” if they do not fit in any of the previous categories.

### Integrative analysis of RNA-seq and CUT&RUN

To identify differential peaks correlated with gene expression changes, we adapted some ideas from the intePareto method ^78^. For each gene *g*, we converted its RNA-seq log_2_(FC) to a z-score by scaling the log_2_(FC) to the standard deviation of all fold-changes in the sample using the following formula:

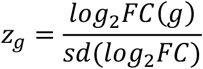

Instead of associating a single peak to each gene as was done in the original intePareto method, we associated a gene to all peaks located within TSS±50kb to be able to unbiasedly identify the most correlated peak. Similarly, we converted the fold change value of each peak *p* in *TSS*_*g*_±50kb to a z-score using the same formula but using ChIP-seq fold change values:

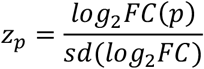

For each gene-peak pair, we calculated a combined z-score by multiplying their z-scores as follows (Fig 5a):

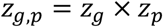

The multi-objective Pareto optimization was then calculated using the ‘psel’ function from the ‘rPref’ R/package (v1.3) [https://doi.org/10.32614/RJ-2016-054]. The peaks from the top 10 best Pareto levels were selected as the most correlated/anti-correlated.

### Statistics and Reproducibility

Statistical analyses were performed using R 4.0.1 or Prism 9.0.2 (GraphPad Software). Parameters of statistical analyses such as the number of replicates and/or experiments (*n*), deviations, p-values, and types of statistics tests are included in the figures or figure legends. For all the *in vivo* experiments, at least three biological replicates were assessed. All *in vitro* assays were performed with at least two independent sample sets. Error bars on graphs represent the mean ±SEM.

## SUPPLEMENTARY DATA

**Supplementary Fig. 1.**
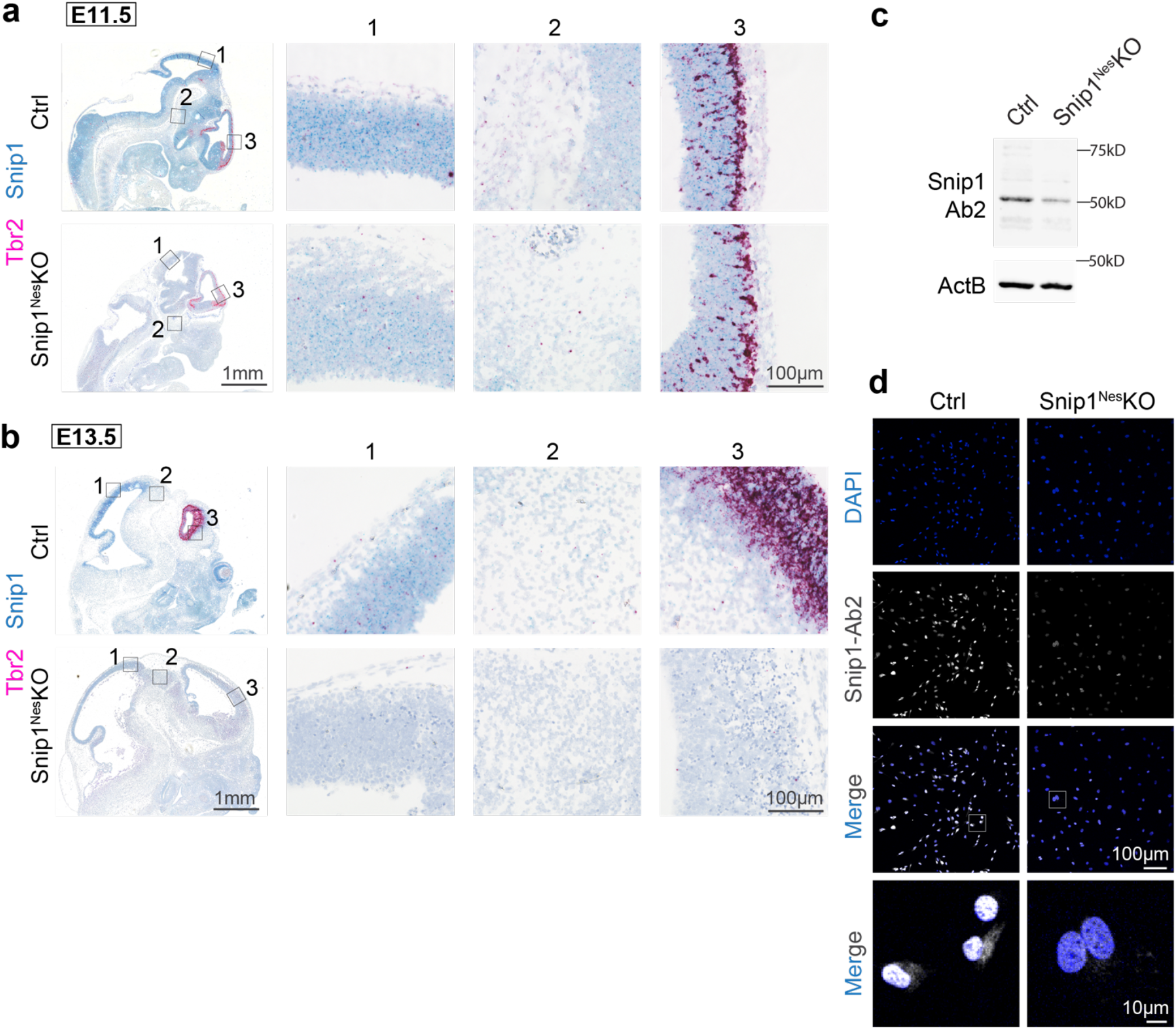
*Snip1* transcript is detected throughout the developing brain with strong expression in the neuroepithelia. **a-b** RNAscope *in situ* hybridization of control and *Snip1*^*Nes*^-KO cryosections at (**a**) E11.5 and (**b**) E13.5. Three magnified representative regions are shown in (**1-3**). At E13.5, robust Snip1 expression (in teal) was detected in the neuroepithelia of the control embryo, whereas it is reduced in the *Snip1*^*Nes*^-KO embryo. Cells positive for the intermediate progenitor marker Tbr2 (in red) were reduced in the *Snip1*^*Nes*^-KO embryo. Bar, 1 mm (entire brain view) and 100 μm (magnified view). **c** WB of control and *Snip1*^*Nes*^-KO brains at E13.5. Snip1-Ab2; anti-Snip1 antibody from ThermoFisher. **d** IF of Snip1 (Ab2; ThermoFisher antibody) and DAPI in control and *Snip1*^*Nes*^-KO cultured NPCs. Bar, 100 μm (entire view) and 10 μm (magnified view).

**Supplementary Fig. 2.**
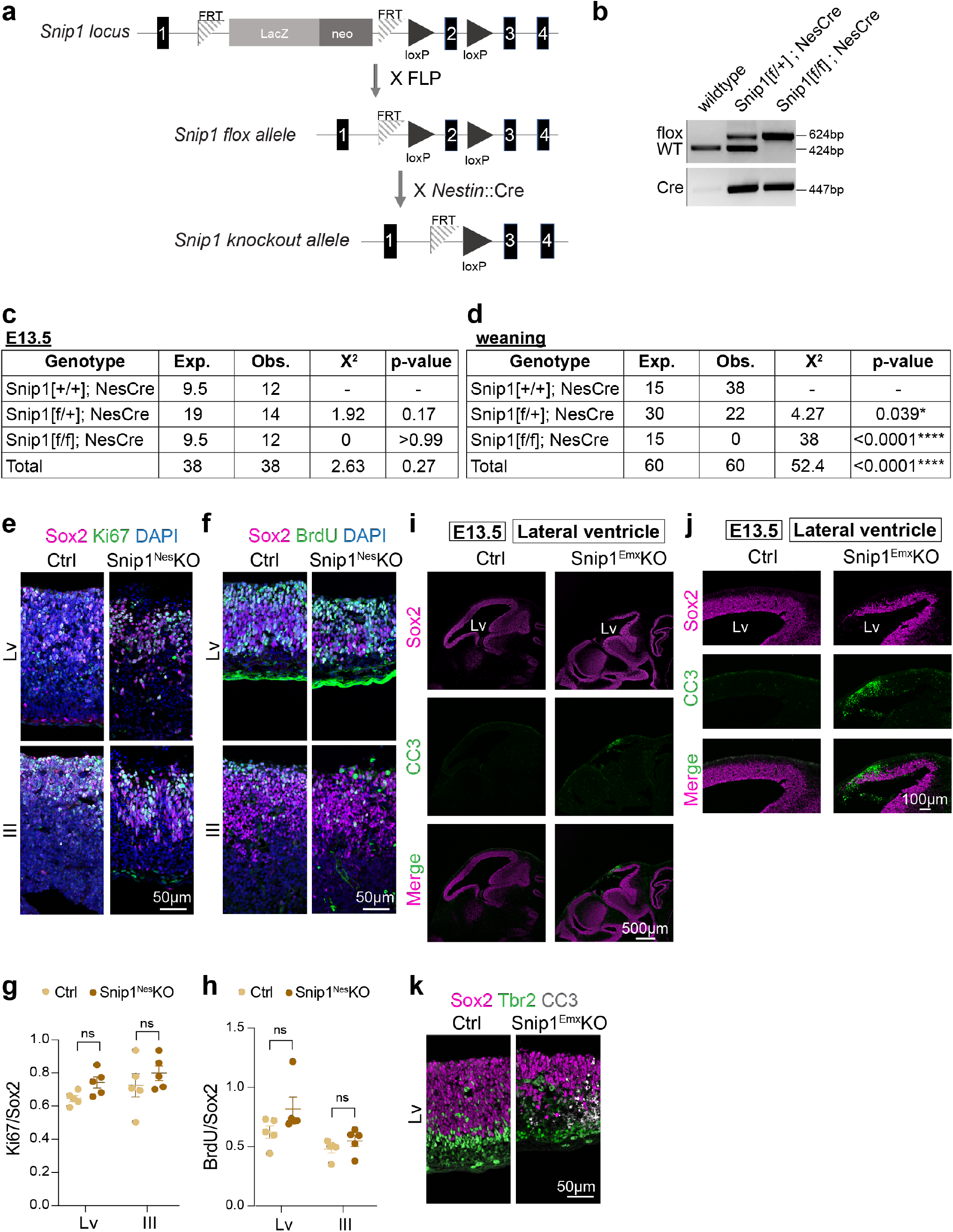
Mice with Snip1-depleted NPCs are not viable by weaning age. **a** Schematic representation of *Snip1* locus containing the LacZ cassette and loxP sites flanking exon 2. Excision of the LacZ cassette by flippase (FLP) generates *Snip1*[flox] allele. Subsequent excision of exon 2 by Cre recombinase driven by *Nestin* promoter (*Nestin*::Cre) generates *Snip1* knockout allele. **b** Genotyping PCR of wildtype allele, *Snip1*[flox] allele, and allele with Cre recombinase transgene. **c-d** Viability of Snip1[+/+]^Nes^, Snip1[flox/+]^Nes^ and Snip1[flox/flox]^Nes^ mice at (**c**) E13.5 and at (**d**) the weaning age (between 3 and 4 weeks after birth). Chi-square test was performed to determine if the mice of three genotypes are viable at the Mendelian ratio. *p <0.05 and ****p <0.0001. *Snip1*^*Nes*^*-*KO embryos were obtained at the expected Mendelian frequency at E13.5; however, no *Snip1*^*Nes*^*-*KO mice were obtained at the weaning age. **e-f** IF of Sox2 and (**e**) Ki67 or (**f**) BrdU in DAPI-stained sagittal cryosections of the E13.5 brain. Germinal zones around the lateral and 3^rd^ ventricles were examined. Bar, 50 μm. **g-h** Quantification of the proliferating NPCs in control and *Snip1*^*Nes*^-KO embryos at E13.5. The plot compares one representative sibling pair and each data point represents one image. Data are presented as mean ± SEM, and two-way ANOVA was used for statistical analysis. ns = not statistically significant. **i-j** IF of Sox2 and CC3 in sagittal cryosections of control and *Snip1*^*Emx*^-KO brains at E13.5. Germinal zones around lateral ventricle was examined. Bar, (**f**) 500 μm and (**g**) 100 μm. **k** IF of Sox2 and CC3 overlayed with an intermediate progenitor marker Tbr2 in control and *Snip1*^*Emx*^-KO brains at E13.5. Bar, 50 μm.

**Supplementary Fig. 3.**
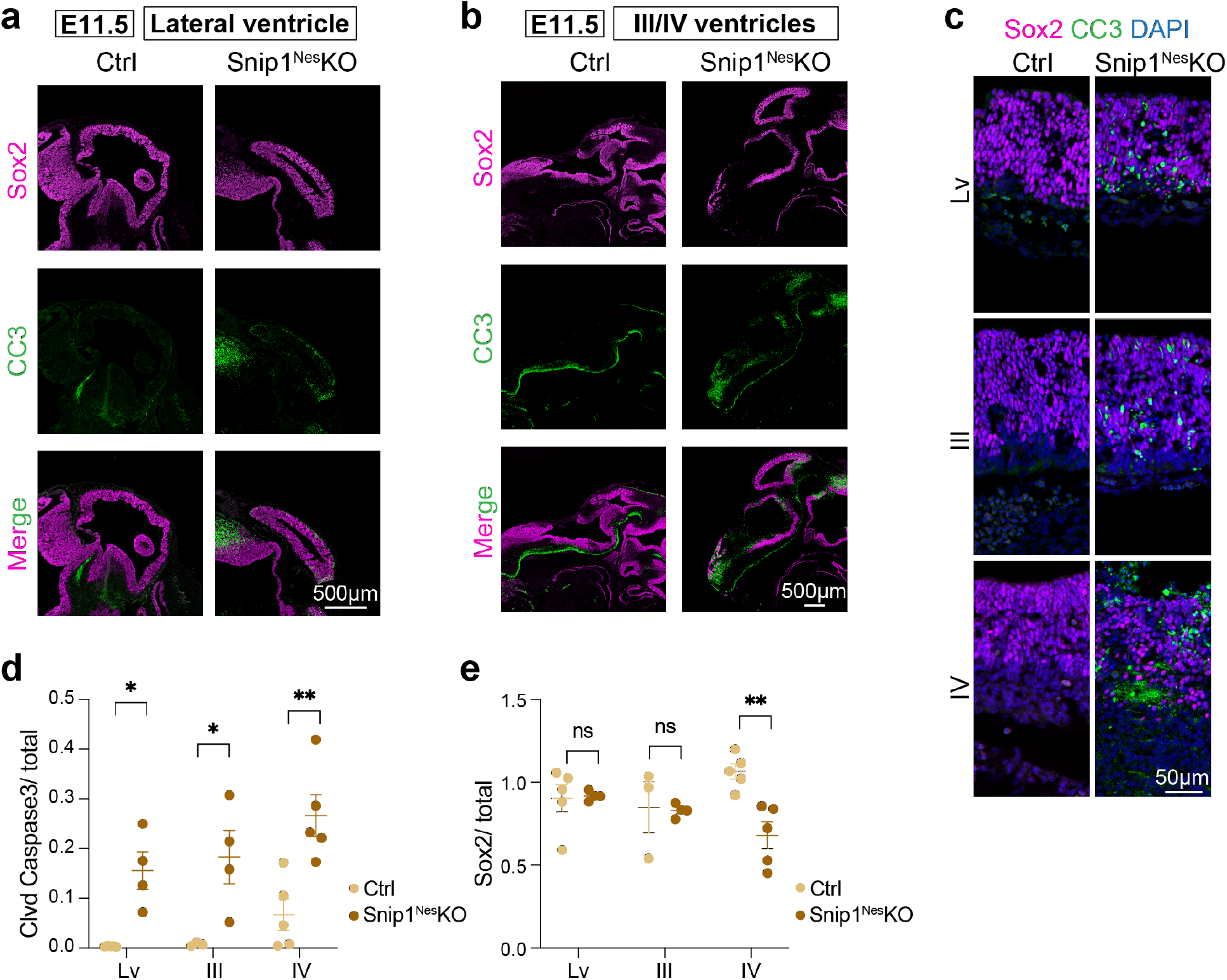
Induction of apoptosis occurs in the *Snip1*^*Nes*^-KO brains as early as E11.5. **a-b** IF of Sox2 and CC3 in sagittal cryosections of the E11.5 brain. Germinal zones around (**a**) lateral ventricle and (**b**) 3^rd^/4^th^ ventricles were examined. Bar, 500 μm. **c** IF with a higher magnification staining against Sox2, CC3, and DAPI of the E11.5 brain. Bar, 50 μm. **d-e** Quantification of CC3-positive and Sox2-positive cells in the neuroepithelial lining of the ventricles of control and *Snip1*^*Nes*^-KO embryos at E11.5. DAPI staining was used to count the total number of cells. Each data point represents one image. Data are presented as mean ± SEM, and two-way ANOVA was used for statistical analysis. ns = not statistically significant; *p <0.05; **p <0.01.

**Supplementary Fig. 4.**
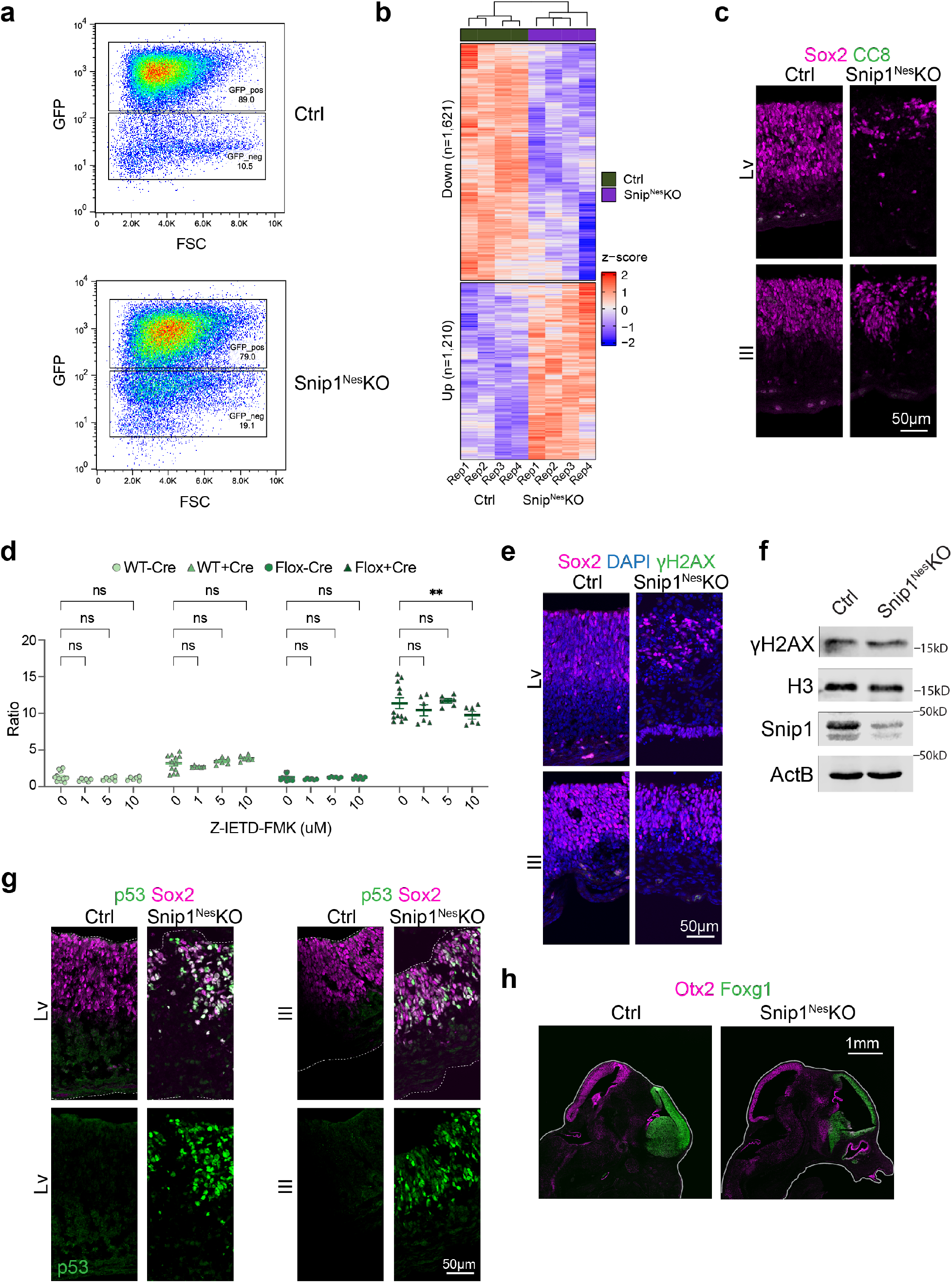
*Snip1*^*Nes*^-KO-induced cell death does not involve DNA damage response. **a** FACS density plots of the control and *Snip1*^*Nes*^-KO brain cells at E13.5. The proportion of the GFP-positive population is reduced in the *Snip1*^*Nes*^-KO brains vs. control brains. **b** Heatmap of differentially expressed genes (by the criterion of FDR<0.05) in *Snip1*^*Nes*^-KO and control NPCs. **c** IF of cleaved caspase 8 (CC8) and Sox2 in sagittal cryosections of E13.5 brains. Bar, 50 μm. **d** Quantification of cells with active caspase 3 quantified by FACS. Snip1 was depleted in Snip1[flox/flox] NPCs by lentiviral mCherry-Cre while cells were treated with a caspase 8 inhibitor Z-IETD-FMK at different concentrations. **e** IF of γH2AX and Sox2 in sagittal cryosections of E13.5 brains. Bar, 50 μm. **f** WB detecting γH2AX in the nuclear extract of control and *Snip1*^*Nes*^-KO brains at E13.5. **g** IF of p53 and Sox2 in sagittal cryosections of E13.5 brains. Bar, 50 μm. Little p53 signal was detected in the control brain, whereas p53 was detected in a large proportion of the Sox2-positive cells in the *Snip1*^*Nes*^-KO brains. **h** IF of Foxg1 (a forebrain marker) and Otx2 (a mid/hindbrain marker) in control and *Snip1*^*Nes*^-KO brains at E13.5. Bar, 1 mm.

**Supplementary Fig. 5.**
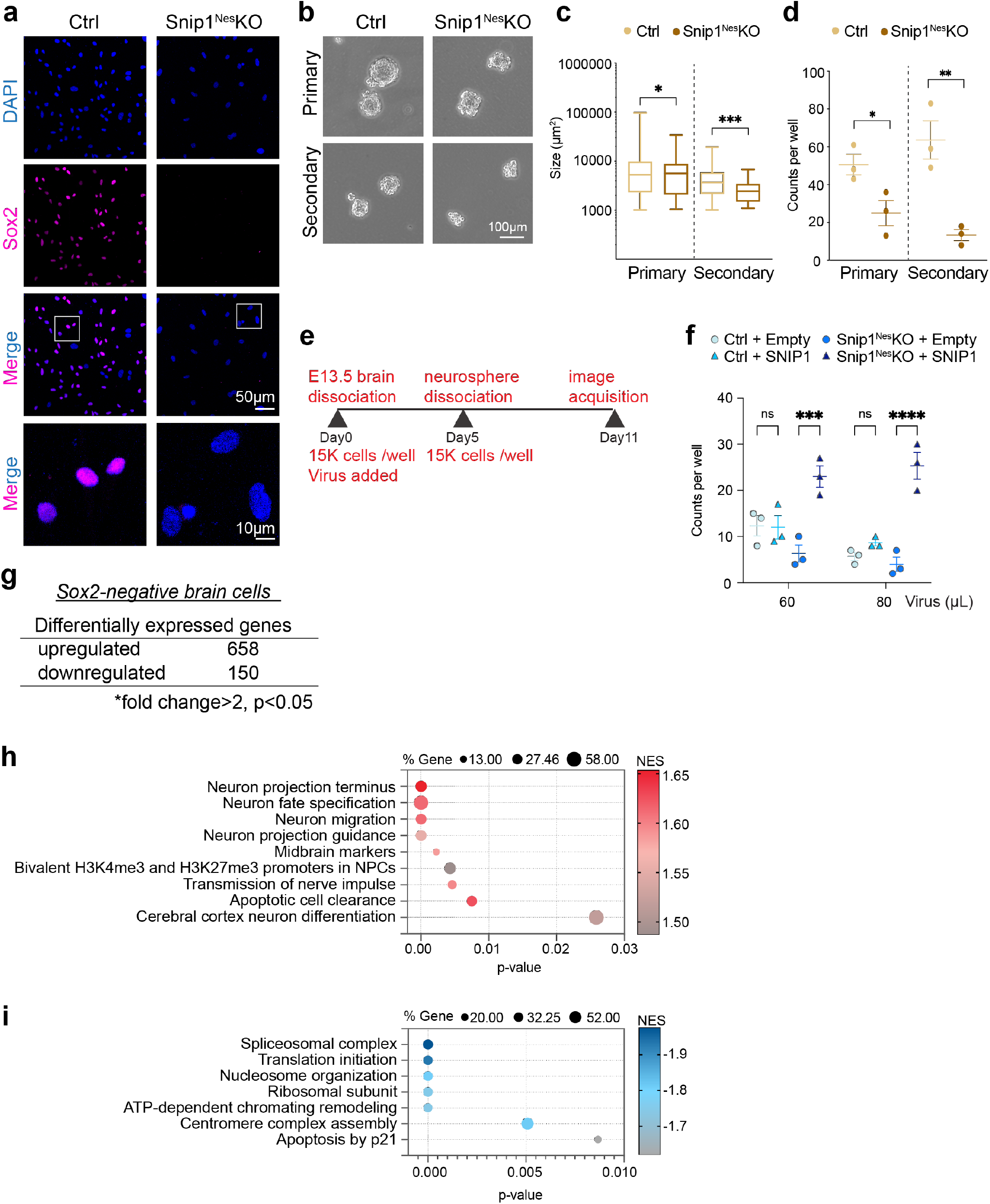
Effect of Snip1 on NPC self-renewal and differentiation. **a** IF of Sox2 and DAPI in control and *Snip1*^*Nes*^-KO cultured NPCs. Bar, 50 μm (entire view) and 10 μm (magnified view). **b** Brightfield images of the sequential neurosphere assay with control and *Snip1*^*Nes*^-KO NPCs. For primary neurosphere formation, 10,000 cells were seeded to each well of low-attachment 6-well plates. For secondary neurosphere formation, primary neurospheres were dissociated and 3,000 cells were seeded to each well. *Snip1*^*Nes*^-KO neurospheres did not grow well and therefore, we failed to obtain the seeding number of 10,000 cells for the secondary neurosphere. All neurospheres were imaged after 5 days of culture. Bar, 100 μm. **c-d** Quantification of the neurosphere size and counts per well. Data are presented as (**c**) box plots and (**d**) mean ± SEM and unpaired t-test was used for statistical analysis. *p <0.05; **p <0.01; ***p <0.001. Snip1-depleted NPCs cannot maintain their self-renewal property in culture. **e** Schematic of a neurosphere rescue experiment. **f** Quantification of the counts of neurospheres per well. Data are presented as mean ± SEM, and two-way ANOVA was used for statistical analysis. ns = not statistically significant; ***p <0.001; ****p <0.0001. Overexpressing human SNIP1 in *Snip1*^*Nes*^-KO NPCs increased neurosphere formation. **g** The numbers of the differentially expressed genes in the Sox2-negative cells of the control and *Snip1*^*Nes*^-KO brains at E13.5. Two RNA-seq datasets each of control and *Snip1*^*Nes*^-KO cells were analyzed. The numbers of genes that passed the cutoff of fold-change >2 and p <0.05 are shown. **h-i** Bubble plots of the enriched gene sets in (**h**) upregulated genes and (**i**) downregulated genes in *Snip1*^*Nes*^-KO vs. control Sox2-negative brain cells. Differentially expressed genes were first ranked by their fold-change and p-value before GSEA was performed.

**Supplementary Fig. 6.**
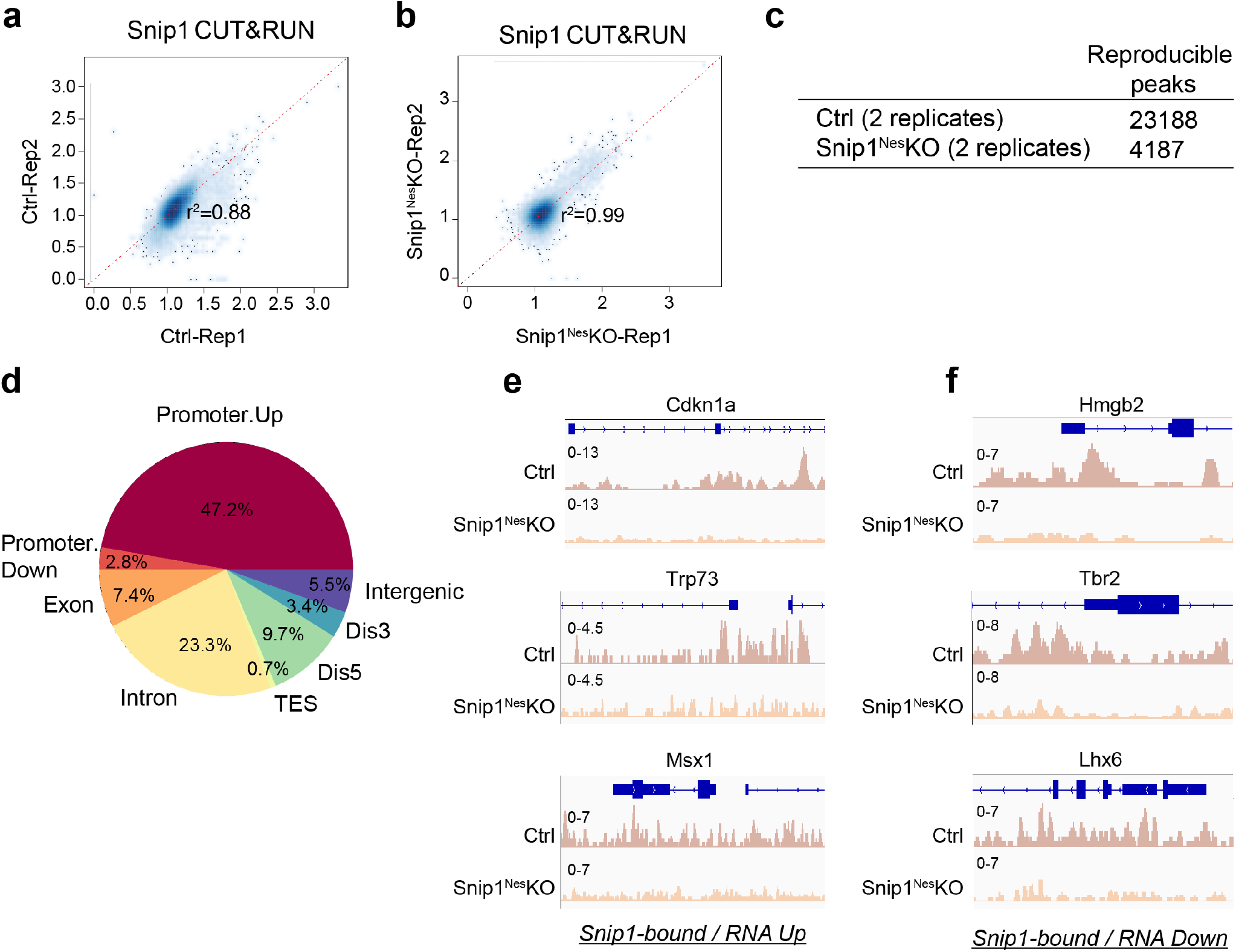
Profiling of Snip1 binding to chromatin by CUT&RUN. **a-b** Pearson correlation plot showing the variance between replicates for Snip1 CUT&RUN. **c** The number of reproducible peaks bound by Snip1 in control and *Snip1*^*Nes*^*-*KO NPCs. Peaks were called by merging both SICER and mac2 peaks with the cutoff of FDR <0.05 and FDR < 0.5. A peak is considered reproducible if it is detected with at least FDR<0.05 in one replicate and no more than FDR>0.5 in the others (see Methods). **d** Proportions of Snip1 binding to genomic features. Fifty percent of Snip1 binding was detected in promoter regions. **e-f** Snip1 CUT&RUN tracks visualized by IGV at (**e**) upregulated genes and (**f**) downregulated genes. *Cdkn1a*, Chr17: 29,090,888 - 29,095,850. *Trp73*, Chr4: 154,132,565 - 154,143,373. *Msx1*, Chr5: 37,818,429 - 37,828,924. *Nes*, Chr3: 87,970,718 - 87,974,908. *Tbr2*, Chr9: 118,476,575 - 118,480,298. *Lhx6*, Chr2: 36,101,041 - 36,106,574.

**Supplementary Fig. 7.**
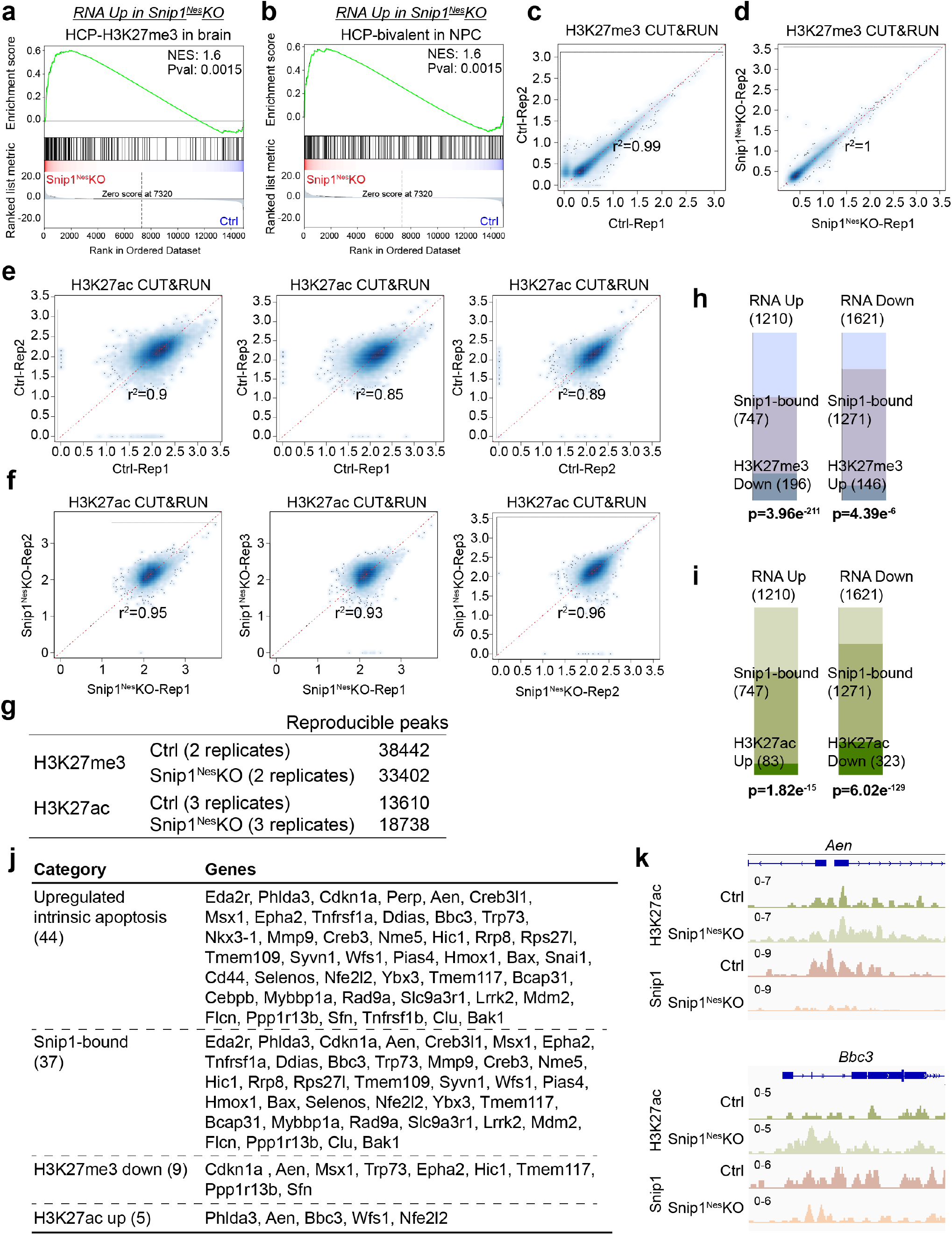
Profiling of H3K27me3 and H3K27me3 in control and *Snip1*^*Nes*^-KO NPCs. **a-b** Representatives of GSEA of upregulated genes in *Snip1*^*Nes*^-KO NPCs. Upregulated genes were enriched in (**a**) high-CpG-density promoters with H3K27me3 in the embryonic murine brain and (**b**) high-CpG-density promoters with bivalent (H3K27me3 and H3K4me3) marks in mouse NPCs. Differentially expressed genes were first ranked by their fold-change and p-value before GSEA was performed. **c-f** Pearson correlation plots between replicates in control and *Snip1*^*Nes*^*-*KO samples for (**c-d**) H3K27me3 and (**e-f**) H3K27ac CUT&RUN. **g** The number of reproducible peaks enriched with H3K27me3 or H3K27ac in control and *Snip1*^*Nes*^*-*KO NPCs. Peaks were called by merging both SICER and macs2 peaks with the cutoff of FDR <0.05 and FDR < 0.5. A peak is considered reproducible if it is detected with at least FDR<0.05 in one replicate and no more than FDR>0.5 in the others (see Methods). **h-i** Bar charts displaying the proportions of differentially expressed genes that were occupied by Snip1 and/or exhibited changes in (**h**) H3K27me3 or (**i**) H3K27ac levels. Hypergeometric test was performed for statistical analysis. **j** Lists of upregulated intrinsic apoptosis genes in *Snip1*^*Nes*^-KO vs. control NPCs under three different categories corresponding to **Fig 3g**. **k** H3K27ac and Snip1 CUT&RUN tracks visualized by Integrative Genomics Viewer (IGV) at upregulated intrinsic apoptosis genes. H3K27ac levels increased at the presented loci in *Snip1*^*Nes*^-KO. *Aen*, Chr7:78,894,346 - 78,897,964. *Bbc3*, Chr7:16,307,660 - 16,311,277.

**Supplementary Fig. 8.**
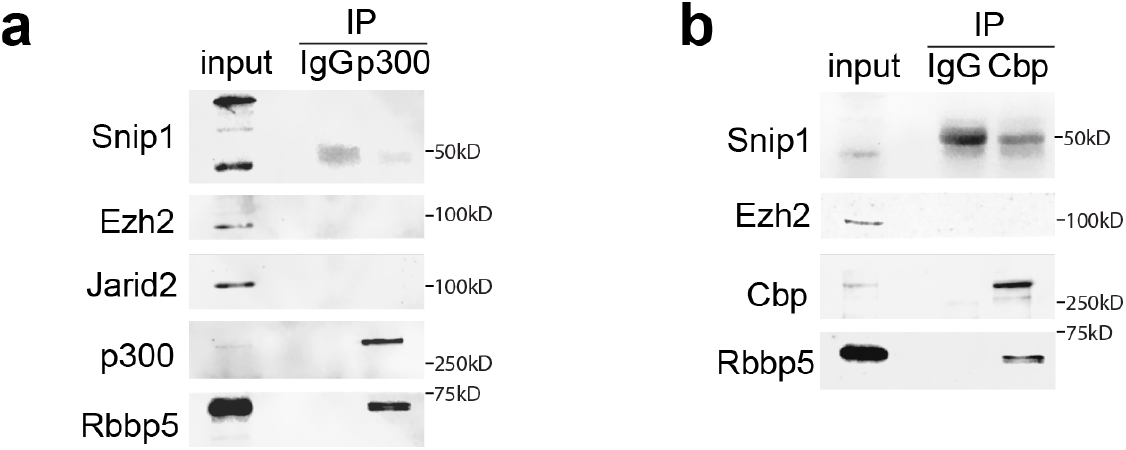
The interactions between Snip1 and p300/Cbp are not detected in NPCs. **a-b** Co-immunoprecipitation followed by WB to examine the interaction between Snip1 and p300/Cbp. (**a**) p300 or (**b**) Cbp was immunoprecipitated in the NPC nuclear extract. Anti-IgG antibody was used as the control for IP and Rbbp5 was a positive control for p300 and Cbp interactions.

**Supplementary Fig. 9.**
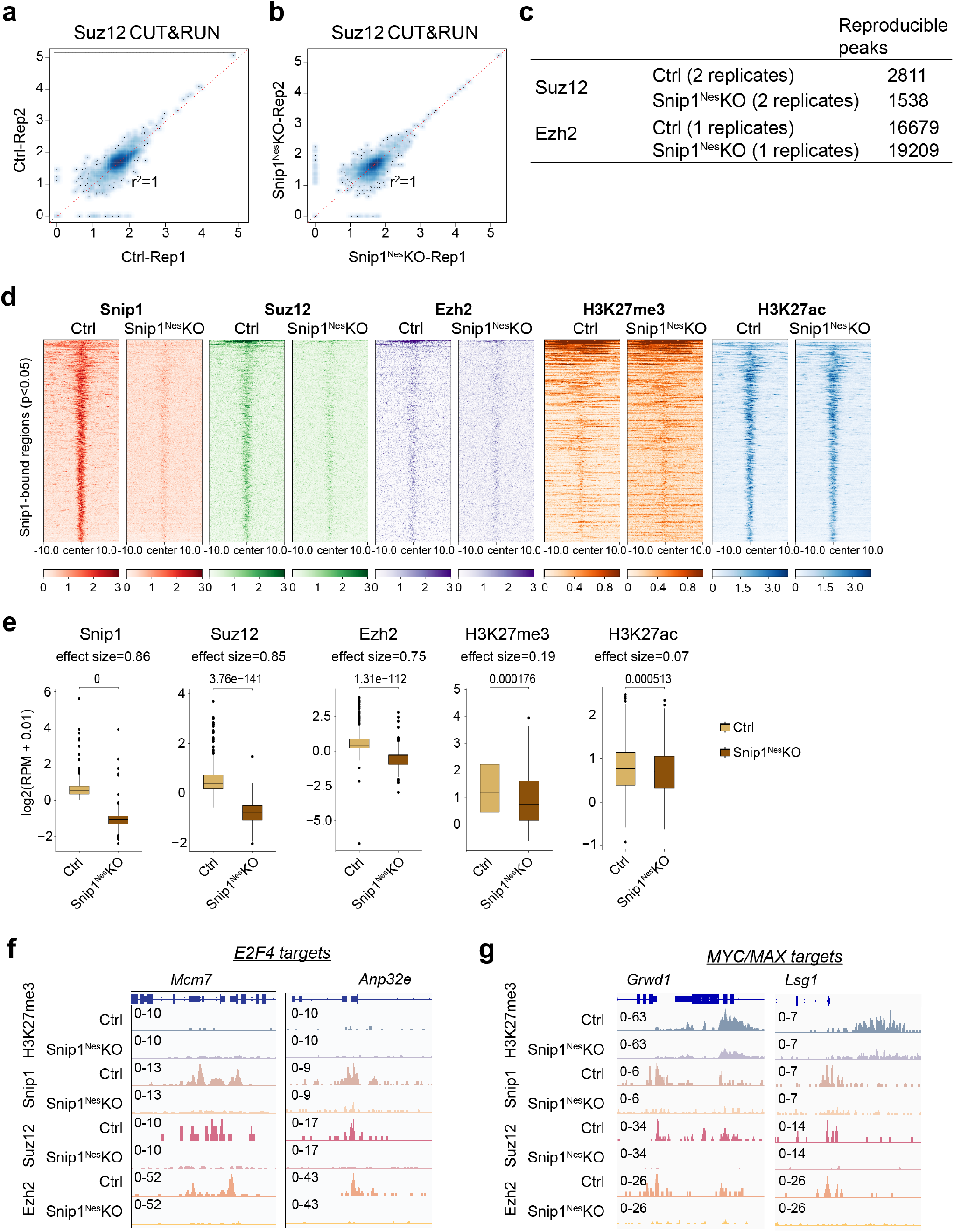
Profiling of PRC2 subunits on chromatin by CUT&RUN. **a-b** Pearson correlation plots between replicates in (**a**) control and (**b**) *Snip1*^*Nes*^*-*KO samples for Suz12 CUT&RUN. **c** The number of reproducible peaks enriched with Suz12 or Ezh2 in control and *Snip1*^*Nes*^*-*KO NPCs. Peaks were called by merging both SICER and macs2 peaks with the cutoff of FDR <0.05 and FDR < 0.5. A peak is considered reproducible if it is detected with at least FDR<0.05 in one replicate and no more than FDR>0.5 in the others (see Methods). **d** Heatmaps aligning peaks of Suz12, Ezh12, H3K27me3, or H3K27ac at the Snip1 targets. Snip1 peaks that were reduced with p <0.05 in *Snip1*^*Nes*^*-*KO vs. control were considered as true Snip1 targets. A dark color indicates high intensity and a light color indicates low intensity. **e** Box plots comparing the binding intensity of Snip1, PRC2, and H3K27me3/ac at the Snip1targets in *Snip1*^*Nes*^-KO vs. control NPCs. **f-g** H3K27me3, Snip1, and PRC2 CUT&RUN tracks visualized by IGV at the reported targets of (**f**) E2F4 (*Mcm7* and *Anp32e*) and (**g**) MYC/MAX (*Grwd1* and *Lsg1*). *Mcm7*, Chr5: 138,169,085 - 138,173,262. *Anp32e*, Chr3: 95,925,524 - 95,933,911. *Grwd1*, Chr7: 45,829,003 - 45,836,327. *Lsg1*, Chr16: 30,584,062 - 30,593,010.

**Supplementary Fig. 10.**
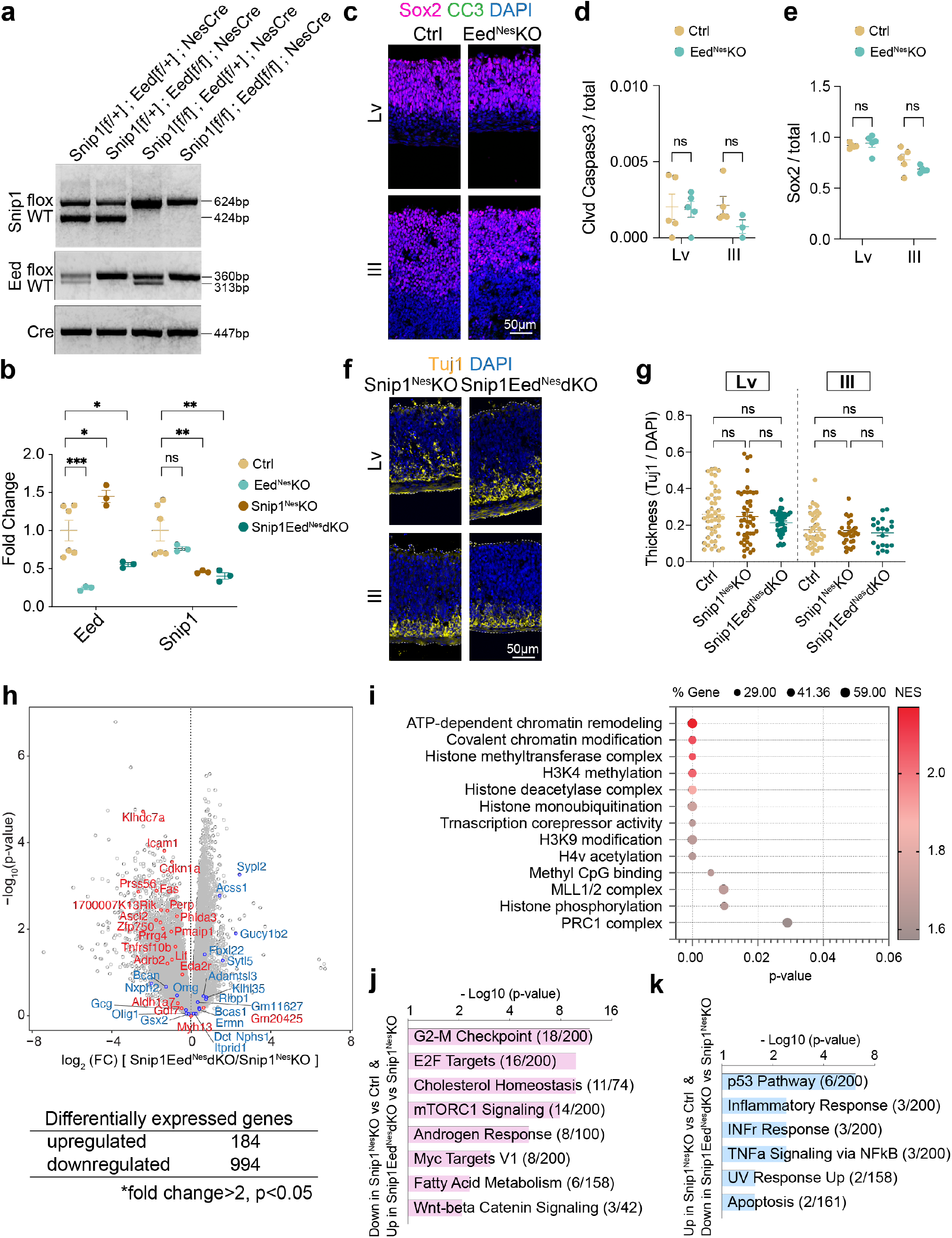
Characterization of *Eed*^*Nes*^*-*KO and *Snip1*^*Nes*^*-Eed*^*Nes*^*-*dKO. **a** Genotyping PCR of *Snip1*-flox allele, *Eed*-flox allele, and allele with *Nes*::Cre recombinase. **b** Quantitative PCR of *Eed* and *Snip1* transcripts in the NPCs from E13.5 brains. The Cq values of each gene were normalized to that of a housekeeping gene *Gapdh*. The expression level of each gene is relative to the level in the control brain. Data are presented as mean ± SEM, and two-way ANOVA was used for statistical analysis. ns = not statistically significant; *p <0.05; **p <0.01; ***p <0.001. **c** IF analysis of Sox2 and CC3 in E13.5 brain. Bar, 50 μm. **d-e** Quantification of CC3-positive and Sox2-positive cells in the neuroepithelial lining of the ventricles of control and *Eed*^*Nes*^-KO embryos at E13.5. DAPI staining was used to quantify the total number of cells. Each data point represents one image. Data are presented as mean ± SEM, and two-way ANOVA was used for statistical analysis. ns = not statistically significant. **f** IF of Tuj1 in the E13.5 brain. Bar, 50 μm. **g** Thickness of the Tuj1-positive region relative to the cortex. Each data point represents one image. N=11 embryos for control, n=7-11 for *Snip1*^*Nes*^-KO, and n=5-7 for *Snip1*^*Nes*^*-Eed*^*Nes*^-dKO. Data are presented as mean ± SEM, and two-way ANOVA was used for statistical analysis. ns = not statistically significant. **h** Volcano plot and the numbers of differentially expressed genes in *Snip1*^*Nes*^*-Eed*^*Nes*^-dKO vs. *Snip1*^*Nes*^-KO. Genes that passed the cutoff of fold-change >2 and p <0.05 were counted. **i** Chromatin modification-related gene sets were enriched in the upregulated genes in *Snip1*^*Nes*^*-Eed*^*Nes*^-dKO vs. *Snip1*^*Nes*^-KO. **j-k** Gene ontology of the rescued genes corresponding to **Fig 5m-n**. Genes were searched against Molecular Signatures Database (MSigDB) Hallmark 2020 using Enrichr ^48^.

**Supplementary Table 1.** List of antibodies.

| Target | Vendor | Catalog # | Assay (dilution) |
| --- | --- | --- | --- |
| Normal Rabbit IgG | RD Systems | AB-105-C | IP<br>CUT&RUN |
| Normal Goat IgG | RD Systems | AB-108-C | IP |
| Jarid2 | Novus Biological | NB100-2214 | IP<br>WB (1:1000) |
| Ezh2 | Active Motif | 39934 | WB (1:1000) |
| Ezh2 | Active Motif | 39076 | IP<br>WB (1:1000)<br>CUT&RUN |
| Suz12 | Cell Signaling Technology | 3737 | WB (1:1000)<br>CUT&RUN |
| Suz12 | Active Motif | 39057 | WB (1:1000)<br>CUT&RUN |
| Snip1 | ThermoFisher | 29412 | IP<br>IF (1:50)<br>WB (1:1000)<br>CUT&RUN |
| Snip1 | ProteinTech | 14950-I-AP | IP<br>WB (1:1000) |
| Rbbp5 | Bethyl Laboratories | A300-109A | WB (1:1000) |
| p300 | RD Systems | AF3789 | IP<br>WB (1:1000) |
| Cbp | GeneTex | GTX101249 | IP<br>WB (1:1000) |
| *Drosophila* H2Av (spike-in control) | Active Motif | 61686 | CUT&RUN (0.25ug) |
| p53 | Leica Biosystems | NCL-L-p53-CM5p | IF (1:100) |
| Histone H3 | Rockland | 100-401-E81 | WB (1:2000) |
| H3K27me3 | Millipore | 07-449 | CUT&RUN |
| H3K27ac | Abcam | ab4729 | CUT&RUN |
| γH2AX | Cell Signaling Technology | 9718 | IF (1:100) |
| γH2AX | Millipore | 05-636 | WB (1:1000) |
| β-Actin | Sigma-Aldrich | A1978 | WB (1:2000) |
| Sox2 | Santa Cruz Biotechnology | sc-17320 | IF (1:150) |
| Tbr2 | ThermoFisher | 14-4875-80 | IF (1:200) |
| Insm1-Alexa Fluor® 488 | Santa Cruz Biotechnology | sc-271408 AF488 | IF (1:50) |
| Tuj1 | Sigma-Aldrich | T8660 | IF (1:300) |
| Map2 | Abcam | ab5392 | IF (1:5000) |
| cleaved caspase 3 (Asp175) | Cell Signaling Technology | 9661 | IF (1:200) |
| cleaved caspase 8 (Asp387) | Cell Signaling Technology | 8592 | IF (1:200) |
| cleaved caspase 9 (Asp353) | Cell Signaling Technology | 9509 | IF (1:200) |
| Ki-67 | Cell Signaling Technology | 9129 | IF (1:100) |
| BrdU | Santa Cruz Biotechnology | 32323 | IF (1:100) |
| Foxg1 | Abcam | ab196868 | IF (1:200) |
| Otx2 | R&D Systems | AF1979 | IF (1:150) |

**Supplementary Table 2.**
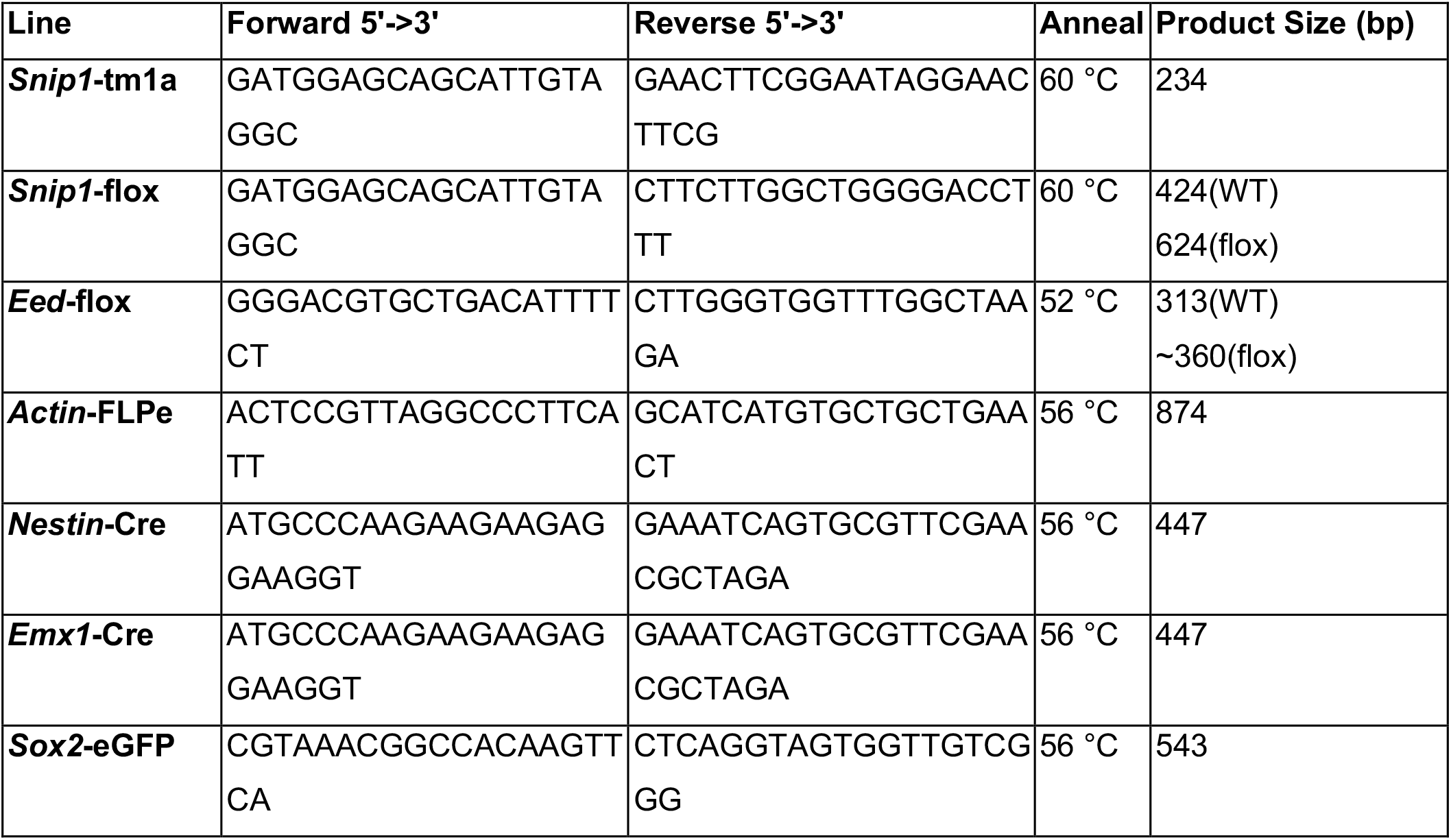
Genotype primers and conditions.

**Supplementary Table 3.**
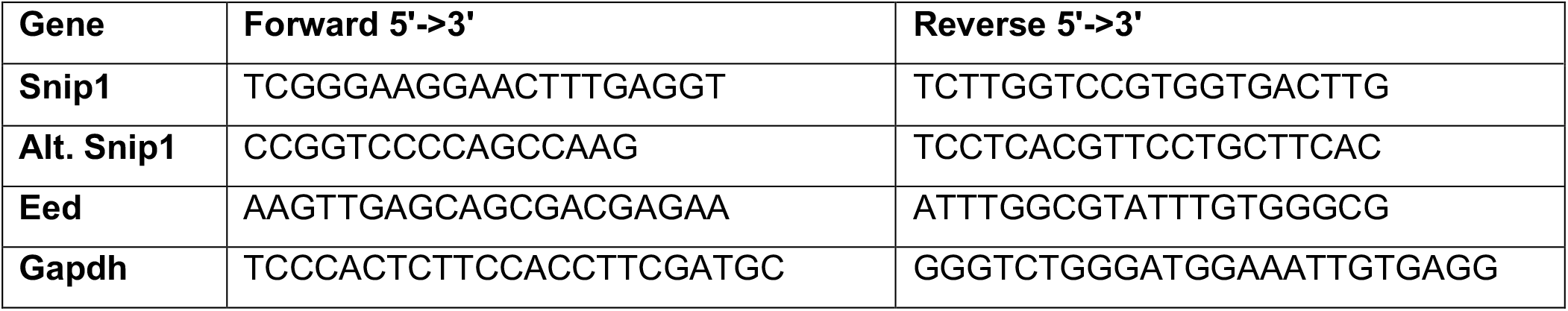
Primers for RT-qPCR.

